# Structural and Dynamic Basis of DNA Capture and Translocation by Mitochondrial Twinkle Helicase

**DOI:** 10.1101/2022.08.11.503323

**Authors:** Zhuo Li, Parminder Kaur, Chen-Yu Lo, Neil Chopra, Jamie Smith, Hong Wang, Yang Gao

**Affiliations:** BioSciences Department, Rice University, Houston, Texas, 77005, USA; Physics Department, North Carolina State University, Raleigh, North Carolina, 27695, USA; Center for Human Health and the Environment, North Carolina State University, Raleigh, North Carolina, 27695, USA; Toxicology Program, North Carolina State University, Raleigh, North Carolina, 27695, USA

## Abstract

Twinkle is the sole helicase responsible for mitochondrial DNA replication. Twinkle can self-load onto and unwind mitochondrial DNA. Nearly 60 mutations on Twinkle have been linked to human mitochondrial diseases. Using cryo-electron microscopy (cryo-EM) and High-speed atomic force microscope (HS-AFM), we obtained the atomic-resolution structure of a vertebrate Twinkle homolog with DNA and captured in real-time how Twinkle is self-loaded onto DNA. Our data highlight the essential role of the non-catalytic N-terminal domain of Twinkle. The N-terminal domain interacts with the helicase domain to stabilize Twinkle hexamers during translocation and the new domain-domain interface is a hotspot for disease-related mutations. The N-terminal domains can protrude approximately 5 nm to capture nearby DNA and initialize Twinkle loading onto DNA. Moreover, structural analysis and subunit doping experiments suggest that Twinkle hydrolyzes ATP stochastically, which explains the low efficiency of Twinkle DNA unwinding and implicates additional regulations of Twinkle during mitochondrial DNA replication.

## INTRODUCTION

A mitochondrion, the powerhouse of a cell, harbors its own genome, which is replicated and maintained differently from its nucleus counterpart. The circular double-stranded mitochondrial genome contains two separate origins (1,2). The replication of the two DNA strands is asynchronous and doesn’t involve Okazaki fragment synthesis (3,4). The mitochondrial replisome shares a distant homology with the replisomes from bacteria and bacteriophages (2,5). Its core components include DNA polymerase γ (Polγ), Twinkle helicase, and mitochondrial single-stranded DNA binding protein (mtSSB) (6). Without a primase, DNA synthesis on both strands is initialized with RNA transcripts produced by the mitochondrial RNA polymerases (7,8). Mutations and deletions in the mitochondrial genome are correlated with numerous neuromuscular diseases and premature aging (9).

Twinkle is the sole mitochondrial replicative helicase (10,11). The expression level of Twinkle is correlated with the copy number of mitochondrial DNA (12), and the homozygous deletion of Twinkle is embryonically lethal (10). Nearly 60 point mutations on Twinkle have been connected to various human mitochondrial diseases (11,13). Patients with Twinkle mutations exhibit reduced mitochondrial DNA copy numbers or partial deletions of the mitochondrial DNA (9). Moreover, Twinkle also participates in mitochondrial DNA repair (14,15) and degradation (16). Consistent with its diverse functions, Twinkle interacts with a number of proteins involved in mitochondrial DNA replication and repair (6,15–17).

Helicases are chemo-mechanical motors that couple ATP hydrolysis to directional translocation on DNA or RNA (18,19). Based on sequence conservations, helicases are divided into 6 superfamilies (SFs), with SF1 and SF2 helicases being monomeric and SF 3-6 helicases being hexameric (19). Replicative helicases are all hexameric and belong to SF3, 4, and 6. The six helicase subunits form a ring- or lock-washer-shaped hexamer, binding ATP at each of the subunit interfaces and DNA in its central channel. DNA binding loops from different subunits stagger like a staircase to interact with the DNA backbone. While hexameric helicases may display different directionality, step sizes, and other mechanistic details, a sequential hand-over-hand translocation mechanism has been proposed for most of them (20–26). Following ATP hydrolysis and product release at one end of the hexamer, the DNA binding loops or the entire subunit moves from one end of the DNA to the other end to form a new ATP binding site. Sequential hydrolysis along the hexamer leads to uni-directional and processive translocation on DNA. A similar hand-over-hand translocation mechanism has been proposed for many hexameric protein translocases (27). Furthermore, the hexameric rings of replicative helicases pose topological restrictions for their loading onto the DNA substrates. To overcome this challenge, bacterial (SF4), archaeal (SF6), and eukaryotic nucleus (SF6) helicases utilize specialized loaders to crack their rings open (28). In contrast, Twinkle (SF4) and bacteriophage T7 gp4 (SF4) helicases are capable of self-loading onto DNA (29,30), but their mechanisms are poorly understood.

Twinkle is a member of the SF4 family helicases and shares low homology with bacterial DnaB and bacteriophage T7 gp4 (10,31). The N-terminal domain of Twinkle is derived from the TOPPRIM family primase (32). However, key residues in the Twinkle primase active site from vertebrates are mutated, and no primase activity has been detected (11). The helicase domain is located on the C-terminal half of Twinkle. Similar to gp4 and DnaB (20,21), Twinkle forms a homo-hexamer and migrates in the 5**’**-to-3**’** direction on the single-stranded (ss)-DNA (31,33). Interestingly, the reported Twinkle helicase activity is significantly lower than that of DnaB or gp4 (34). In addition, Twinkle can bind to and migrate on double-stranded (ds) and Holliday junction DNA (34,35). To date, structures of Twinkle in the absence of DNA have been reported (33,36–38). However, how Twinkle functions mechanistically on DNA and how disease-related mutations affect Twinkle function are still largely unclear. Here we report cryo-EM, biochemical, and HS-AFM characterization of a vertebrate Twinkle homolog. Our data elucidate unique structural and dynamic features of Twinkle helicase, and support a novel molecular model explaining Twinkle self-loading, DNA unwinding, and human disease-related mutations on Twinkle.

## MATERIAL AND METHODS

### Sequence Analysis

Twinkle homolog sequences from *Arabidopsis thaliana, bacteriophage T7, Dictyostelium discoideum, Dictyostelium purpureum, Drosophila melanogaster, Gallus gallus*, and *Homo sapiens* were used as bits for the BLAST search (https://www.uniprot.org/blast/). The target database was set to be UniProtKB reference proteomes plus Swiss-Prot with the E-Threshold set to be 10. Redundant sequences were eliminated, and sequences with less than 400 residues or more than 800 residues were removed. The remaining sequences were aligned with MUSCLE sequence alignment tool (39). The sequence identity matrix was plotted with Matlab.

### Plasmid construction

The LcTwinkle (Entrez ID 108894827, synthesized by GeneUniversal Inc.) was cloned into a modified pET28a vector with an N-terminal histidine tag and a PreScission protease cleavage site. Mutations of LcTwinkle were generated using methods described in the QuikChange mutagenesis kit (Agilent). Sequences of the LcTwinkle constructs were confirmed by sequencing the entire reading frames of each construct. The active site mutation of Hs (uniport Q86RR1), Mm (uniport Q8CIW5), and Dr (uniport A0A0R4ICC1) Twinkle were all synthesized by GeneUniversal Inc. and cloned into the same pET28a vector.

### Protein expression and purification

The LcTwinkle plasmid was transformed into *E. coli* BL21 (DE3) (Novagen). Isopropyl-β-D-thiogalactopyranoside (1 mM) was added to the cell culture at an optical density of 0.8 for protein induction. The *E. coli* cells were further incubated at 16°C for 16 hours under shaking at 150 rpm for protein expression. The cells were harvested and disrupted by sonication in a Lysis buffer containing 50 mM Tris-HCl, pH 8.0, and 1 M NaCl, and 5% glycerol. The soluble fraction was loaded onto a 5 ml HisTrap column (GE Healthcare), which was preequilibrated with the Lysis buffer plus 20 mM imidazole. The column was subsequently washed with 300 ml Lysis buffer plus 50 mM imidazole, and the protein was eluted with 5 ml Lysis buffer plus 300 mM imidazole. The eluted protein was treated with 10 U PreScission protease (Sigma) at 4°C for 2 hours. Afterward, the protein was diluted into a MonoQ buffer containing 50 mM Tris pH (8.0), 0.1 mM ATP, 1 mM MgCl2, 150 mM KCl, 3 mM dithiothreitol (DTT), 5% glycerol and loaded onto a MonoQ column (GE Healthcare) preequilibrated with the MonoQ buffer. LcTwinkle was eluted by a gradient KCl concentration. The purified LcTwinkle was aliquoted and flash-frozen at −80°C for further study. Mutant LcTwinkle proteins were expressed and purified according to the same protocol. Hs, Mm, and Dr Twinkle were expressed and purified similarly as LcTwinkle. After elusion from HisTrap columns, the proteins were mixed with ssDNA (5’-TGGTCTTTTTTTTTTTTTTTTTTTTTTTTT-3’) at 1:1.5 molar ratio in a buffer containing 50 mM Tris (pH 8.0), 150 mM KCl, 3 mM DTT, 1 mM ATP, and 2 mM MgCl2. Following incubation on ice for 10 min, the LcTwinkle-DNA complex was loaded onto a Superose6 column (GE Healthcare) equilibrated in the same buffer.

### ATPase assay

LcTwinkle ATPase activity was assayed with radioactive ATP (γ-32P) and thin-layer chromatography. All reactions were performed in a reaction buffer containing 25 mM Tris-HCl (pH 8.0), 12 mM MgCl2, 3 mM DTT, 0.05-0.5 μM of WT or mutant LcTwinkle, 1 μM ssDNA (5’-GGATTATTTACATTGGCAGATTCACC-3’), and a desired concentration of ATP (with 0.2 μCi labeled γ-32P-ATP). Reactions were carried out at 20°C for 30 min and terminated with EDTA. 2 μl of terminated reaction mix were spotted onto polyethyl-enimine-cellulose plates (Merck, Germany). ATP and released phosphate were then separated chromatographically in a buffer of 0.5 M LiCl. Plates were exposed to a phosphor screen (GE Healthcare) for 4 h. Phosphor screens were imaged using a Sapphire Biomolecular Imager-RGB IS1025 (Azure BioSystems). The data were analyzed with AzureSpot software equipped in the Sapphire imager. The data were fitted to Mekanies-Menten equation with GraphPad (GraphPad Software LLC). All results were based on at least three independent tests.

### DNA unwinding assay

The helicase reactions were carried out in a buffer containing 50 mM Tris-HCl (pH8.0), 50 mM KCl, 5% glycerol, 0.1 mg/ml BSA, 5 mM MgCl2 and 4 mM ATP. Each reaction contained 25 nM FAM-labeled fork DNA substrate, which was annealed with 5’-FAM-CCTAGCTCAGGTTCAGTACTCGAACTCTACATAACTATACATGAATATCATAACTAATAA-3’ and 5’-TTATTAGTTATGATATTCATGTATAGTTATCATCTCAAGCTCATG-3’. The reactions were all performed at 37°C for 30 min. The reactions were stopped with a quench buffer containing 20 mM EDTA, 1% SDS, and 0.2 mg/ml proteinase K. The substrate and product were separated on a 12% native polyacrylamide gel, and the images were analyzed using a Sapphire Biomolecular Imager-RGB IS1025 (Azure BioSystems). All results were based on at least three independent tests.

### DNA binding assay

The DNA binding assay was performed in 100 μl reaction buffer containing 25 mM Tris, 150 mM KCl, 0.5 mM ATP and 1 mM MgCl2, 20 nM Cy5 labeled ssDNA (5’-Cy5-GGATTATTTACATTGGCAGATTCACC-3’) and varying concentrations of WT or mutant LcTwinkle. The reaction mix was loaded onto 96 well plates and the fluorescence anisotropy was measured with 645 nm excitation and 670 nm emission using Infinite M1000 Pro microplate reader (TECAN). The K_D_ was analyzed similarly as in (34) with GraphPad software (GraphPad Software LLC).

### Subunit doping

The CFP and YFP-Twinkle were purified similarly as WT Twinkle. For the FRET experiments, the individual CFP-Twnk and YFP-Twnk proteins or the mixture of two were prepared in a buffer containing 500 KCl, 50 mM Tris and 3 mM DTT. 1 mM ATP and 2 mM MgCl2 were added when indicated. The protein concentration is 0.2 mg/ml for each of the protein. The protein sample is scanned at room temperature using fluorometer (Cary Eclipse Fluorescence Spectrophotometer, Agilent, Santa Clara, CA, USA). During the scan, a 430 nm filter was used as the excitation wavelength and the emission between 430 and 570 nM were collected. For subunit doping experiments, WT and mutants Twinkle were buffer exchanged to 500 KCl, 50 mM Tris and 3 mM DTT. All mixed proteins were incubated at room temperature for 5 minutes before activity assay. The ATPase, helicase, and DNA binding activities were determined as descripted above.

### Negative staining EM

WT and mutant Twinkle were diluted to 0.02 mg/ml in a buffer containing 50 mM Tris (pH 8.0), 150 mM KCl, 3 mM DTT, 1 mM ATP, and 2 mM MgCl2. The ssDNA substrate (5’-TGGTCTTTTTTTTTTTTTTTTTTTTTTTTT-3’) was added at 1:2 molar ratio relative to the LcTwinkle hexamers. 3 μl of LcTwinkle-DNA sample was deposited onto a freshly glow-discharged CF400-CU carbon grid (EMS Inc.). After absorbing the extra protein solution with filter paper, 3 μl of 2% uranyl acetate were added to the grid. Following 20 s incubation, the extra uranyl acetate was removed with filter paper. The grids were imaged on a JEOL2100 microscope operated at 200 kV voltage and 400,000 magnifications. The CTF estimation and 2D classification were performed with RELION 3.0 (40).

### Cryo-EM sample preparation

E325Q LcTwinkle was mixed with ssDNA (5’-TGGTCTTTTTTTTTTTTTTTTTTTTTTTTT-3’) at 1:1.5 molar ratio in a buffer containing 50 mM Tris (pH 8.0), 150 mM KCl, 3 mM DTT, 1 mM ATP, and 2 mM MgCl2. Following incubation on ice for 10 min, the LcTwinkle-DNA complex was loaded onto a Superose6 column (GE Healthcare) equilibrated in the same buffer. The eluted LcTwinkle-DNA complex was concentrated to 1 μM (~0.4 mg/ml) concentration, as determined by the Bradford assay. 3 μl of LcTwinkle-DNA sample was deposited onto a freshly glow-discharged Quantifoil R1.2/1.3 300 mesh grid and blotted using a Vitrobot (FEI) with the standard Vitrobot filter paper, Ø55/20 mm (Ted Pella). The blotting time was set to 4 s, the blotting force was set to 4 and the blotting was done under 100% humidity at 20°C. The grids were flash-frozen in liquid ethane and stored in liquid nitrogen.

### Cryo-EM data collection and processing

6490 micrographs of LcTwinkle-DNA complexes were collected on a Titan Krios electron microscope operated at 300 kV (cryo-EM core facility at University of Texas McGovern Medical School) using the super-resolution mode with a nominal magnification of 130 K (calibrated pixel size of 1.07 Å on the sample level, corresponding to 0.535 Å in superresolution mode). Movies were recorded with a K2 Summit camera, with the dose rate at the detector set to 7 e^-^.s^-1^.Å^-2^. The total exposure time for each video was 7 s, which was fractionated into 35 frames of sub-images. The defocus values ranged between 0.6 and 3 μm. MotionCor2 (41) was used for drift-correction and electron-dose-weighting for all movies. The defocus values were estimated on non-dose-weighted micrographs with Gctf (42). 1976802 particles were picked from 6334 manually screened micrographs with Gautomatch (Developed by Kai Zhang, http://www.mrc-lmb.cam.ac.uk/kzhang/Gautomatch/). After 2D classification with RELION (40), 939599 particles were selected. An *ab initial* hexamer LcTwinkle model was generated in cryoSPARC (43). The particles were separated into six classes by 3D-classification in RELION (40) and two classes (one hexamer and one heptamer) with clear protein features were selected for subsequent processing. Further 3D-classification were performed with cryoSPARC (43) to clean the datasets and separate different conformations of LcTwinkle. For the LcTwinkle-DNA complex, the particles from cryoSPARC were analyzed with Ctf Refine and polishing in RELION (40). Final refinements were down with the non-uniform refinement in cryoSPARC (43).

### Model building and refinement

A homolog model of LcTwinkle was generated with the I-TASSER server (https://zhanglab.dcmb.med.umich.edu/I-TASSER/) and used for the initial rigid-body search. SsDNA coordinates adapted from gp4-DNA structure (PDB ID: 6N7V) were used as starting model for DNA. Each of the protein chains and the DNA were manually docked into the cryo-EM density maps in Chimera (44). The models were first manually adjusted in COOT (45) and then refined in Phenix (46) with real-space refinement and secondary structure and geometry restraints. Due to the low resolution of the NTDs, poly-alanine models were used during the refinement, excepting the residues at the NTD-CTD interfaces with good sidechain densities. For the lower resolution LcTwinkle-DNA_2_, LcTwinkle_6_, and LcTwinkle_7_ structures, the monomers from the refined LcTwinkle-DNA model were docked into the cryo-EM density with Chimera (44) and the connections were adjusted in COOT (45). Statistics of all cryo-EM data collection and structure refinement are shown in Table S1.

### High-speed atomic force microscopy (HS-AFM) imaging in liquids

LcTwinkle (35 nM) was incubated with the linear DNA substrate containing a 37-nt ssDNA gap positioned at 23% from one DNA end (47) in Twinkle Reaction Buffer (20 mM HEPES pH 7.6, 150 mM NaCl, 7.5 mM MgCl2) containing ATP (4 mM) for 1 min. The sample was diluted in Twinkle Reaction Buffer and deposited onto a freshly prepared 1-(3-Aminopropyl)silatrane (APS)-treated mica surface. The sample was incubated with mica (SPI) for 2 min. The APS-mica surface containing the sample was washed with Twinkle Reaction Buffer and scanned in Twinkle Reaction Buffer using a Cypher VRS AFM (Asylum Research). BlueDrive Photothermal Excitation was used to drive a BioLever fast (AC10DS) cantilever with the resonance frequency (f) at ~1500 kHz and the spring constant (k) at ~0.1 N/m. The images were scanned at 0.8-2 frames/s. The images were analyzed using Asylum software.

## RESULTS

### The overall structure of Twinkle helicase

Previously, poor protein solubility hampered structural studies of Twinkle (33,48). Although a high-resolution structure of *Homo sapiens* (Hs) Twinkle was reported recently (38), it is with a disease-related mutant and the DNA and ATP are absent. We attempted to identify a Twinkle homolog similar to the human Twinkle and amenable to high-resolution structural determination. Sequence alignment suggested that vertebrate Twinkle proteins share high identities and may have similar structures and functional properties (Supplementary Figure 1). We picked Twinkle homologs from *Homo sapiens* (Hs), *Mus musculus* (Mm), *Danio Rerio* (Dr), and *Lates calcarifer* (Lc) for testing their stability and protein-DNA complex formation. LcTwinkle shares 65% identity and 90% similarity with HsTwinkle, with most of the essential residues for Twinkle activities and disease-related mutations conserved (Supplementary Figure 1B). Two variants of LcTwinkle sequences were predicted by different databases, one longer variant and one shorter variant lacking the N-terminal mitochondrial targeting sequence (MTS) and the Zinc-binding domain (ZBD) (Figure 1A and Supplementary Figure 1C). The core regions, including the helicase and primase-like domains, are identical in the two variants. The shorter LcTwnk was chosen for cloning. Hs, Mm, Dr, and LcTwinkle homologs with active site mutations corresponding to gp4 E343Q (49) were preliminary purified with an affinity column. Considering the similarities between gp4 and Twinkle, the same ssDNA substrate as in gp4 structure determination (5’-TGGTCTTTTTTTTTTTTTTTTTTTTTTTTT-3’) (20) was used for Twinkle complex formation, although the primase recognition motif TGGTC turned out to be unnecessary at the end. Notably, among all Twinkle proteins tested, only LcTwinkle migrated as a single sharp peak in gel filtration column in the presence of ATP and ssDNA (Fig. 1B and Supplementary Figure 2).

**Figure 1.**
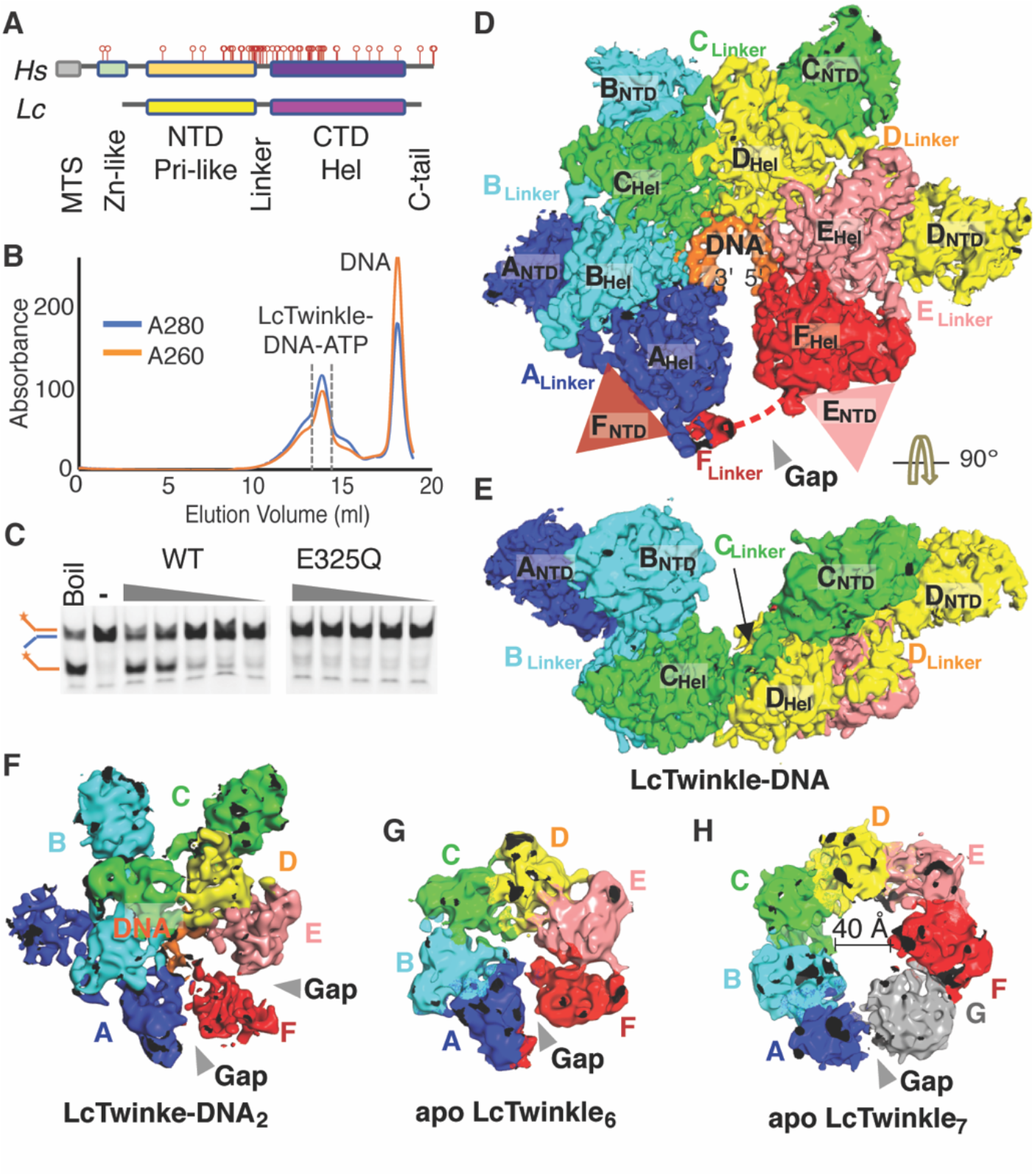
Overall structure of LcTwinkle. **A.** Domain structures of human (*Hs*) and *Lates calcarifer* (*Lc*) Twinkle. The disease-related Twinkle mutations are marked as red bars. MTS: mitochondrial targeting sequence; Zn: Zinc-binding domain; Pri: primase; Hel: helicase; C-tail: C-terminal tail. **B.** E325Q LcTwinkle-DNA complex formation revealed by size exclusion chromatography. **C.** Helicase activity of WT and E325Q LcTwinkle. **D-E.** Two views of the LcTwinkle-DNA complex. The subunits A to F are color-coded from blue to red, following their order on DNA. DNA is colored orange. The NTDs from subunits E and F are disordered and their positions are marked by semi-transparent triangles. The disordered part of the N-C loop in subunit F is indicated by a dashed line. **F.** Structure of LcTwinkle-DNA_2_. The mobile subunit F and the structural gaps between F and its neighboring subunits are highlighted. **G-H.** Structures of apo LcTwinkle hexamer (**G**) and heptamer (**H**). The structural gaps in the oligomers are marked.

Wild-type (WT) and E325Q LcTwinkle were thoroughly purified for further characterization. The purified WT LcTwinkle displayed robust ATP-dependent helicase activity (Fig. 1C). E325Q mutation eliminated the helicase unwinding activity of LcTwinkle (Fig. 1C). Although WT LcTwinkle can bind ssDNA in the presence of non-hydrolyzable ATP analogs (Supplementary Figure 2D), the resulting ssDNA complex was unstable. It migrated as a much higher molecular weight peak in gel filtration (Supplementary Figure 2E). Consistently, the WT LcTwinkle-ssDNA-ATPγS complexes appeared as aggregates on cryo-EM grids (Supplementary Figure 2F). Thus, the purified E325Q LcTwinkle-DNA-ATP complex from gel filtration (Fig. 1B) was used for cryo-EM analysis. We determined the LcTwinkle-DNA complex structure with cryo-EM at 3.5 Å resolution (Fig. 1D, Supplementary Figure 3, Supplementary Table 1). The local resolutions of the C-terminal helicase domains (CTD) were near 3 Å (Supplementary Figure 3E). Atomic structures of all six CTDs can be built (Supplementary Figure 4). The N-terminal primase-like domains (NTD) were at 5-8 Å resolution (Supplementary Figure 3E). The α-helices in the NTDs can be traced, and bulky side chains near the NTD-CTD interfaces are visible (Supplementary Figure 4). The homology model of the LcTwinkle NTD can be confidently docked into the cryo-EM densities in four out of the six subunits (Fig. 1D). The CTD domains of LcTwinkle are similar to the apo structures of HsTwinkle, with a RMSD ~ 1 Å (Supplementary Figure 5A), whereas the NTD domains are less conserved and the RMSD between LcTwinkle and HsTwinkle is ~2.5 Å (Supplementary Figure 5B).

LcTwinkle forms a homo-hexamer wrapping DNA in its central channel (Fig. 1D). The six subunits are labelled as A to F following their order on DNA in the 3’-to-5’ direction. The NTDs are attached to the side of the CTDs. The domain structure of LcTwinkle is consistent with previously published low-resolution structural models of Twinkle (33,36,37), but different from the apo structure of HsTwinkle (38), where the NTD interacts with the CTD from the same subunit (Supplementary Figure 5A). This arrangement of the NTDs is also distinct from all the other replicative helicases, in which the NTDs are all located on top of the helicase rings at the 5’-side of the DNA (Supplementary Figure 5, C-H) (20–23,25). In addition, this structure Compared to gp4, the LcTwinkle NTD swings nearly 90 degrees to the side (Supplementary Figure 5I). Furthermore, the pseudo-primase active site is flipped to face downward, making interactions with the unwound DNA impossible (Supplementary Figure 5I). An extended linker connects the NTD and the CTD (N-C linker) in a domain-swapped manner (Figure 1D, E). The N-C linker from one subunit interacts with the CTD of the second subunit on its 5’-side of DNA. The CTDs form a non-planar lock-washer-shaped hexamer (Figure 1D), similar to gp4 and DnaB (20,21). The subunits at the two ends of the lock-washer are bridged by the N-C linker, making the hexamer a complete circle (Figure 1D). ATPase sites are located at each of the subunit interfaces (Figure 2A), and five ATP molecules can be refined (Supplementary Figure 4E). A 12-nucleotide (nt) ssDNA binds to the central channel of the LcTwinkle, with an average of 2 nucleotides per subunit (Figure 2A and Supplementary Figure 4F).

**Figure 2.**
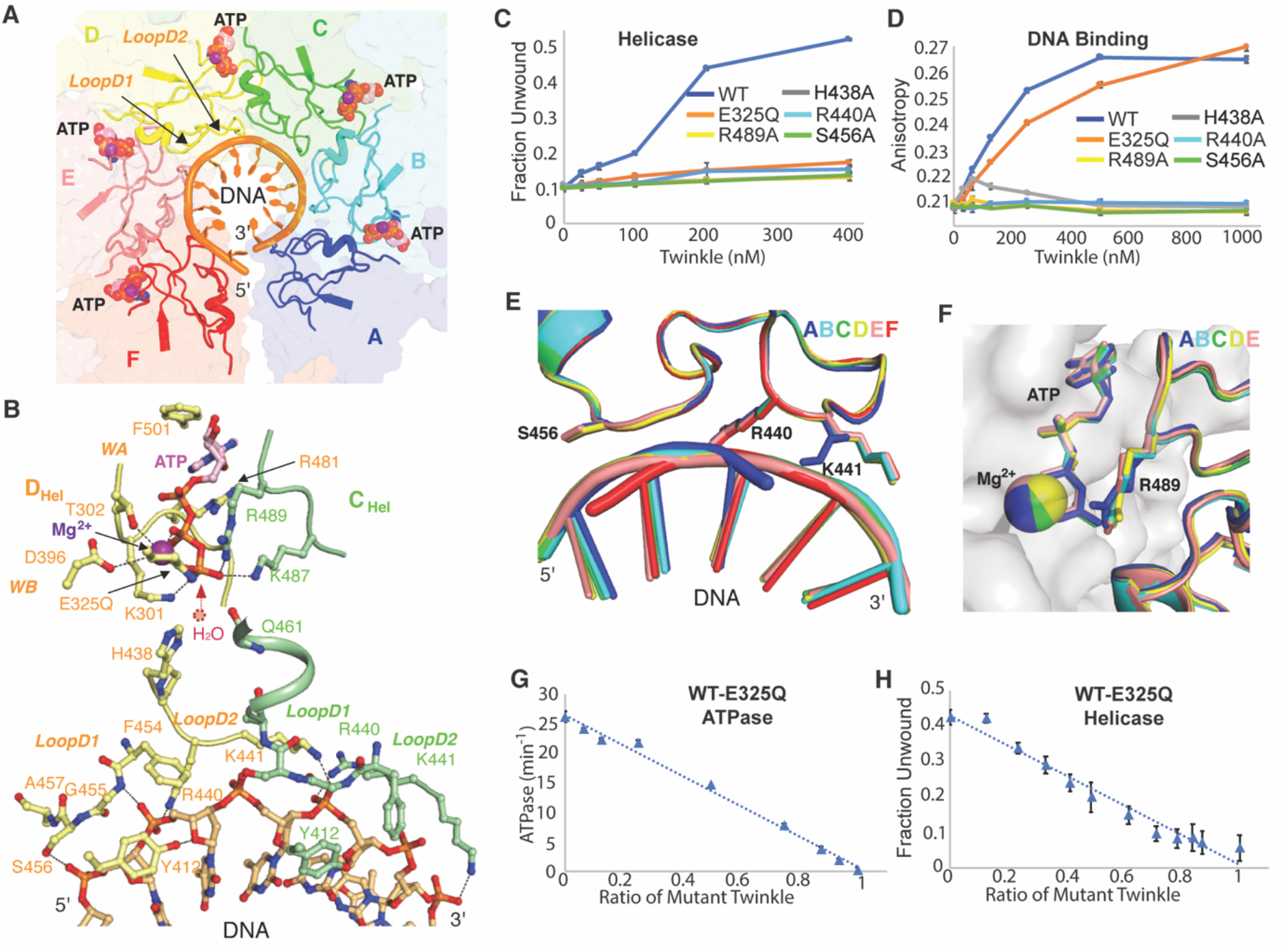
ATP and DNA binding interfaces of LcTwinkle. **A.** ATPase and DNA binding sites in the LcTwinkle-DNA complex. The CTDs are shown as transparent surfaces and structural elements involved in ATP and DNA binding are shown in the cartoon. **B.** Zoom-in view of the ATP and DNA binding sites between subunits C and D. Key residues are shown as sticks. WA, WB, D1, D2 are for Walker A, Walker B, LoopD1, and LoopD2 motifs, respectively. The position of the predicted water molecule for the nucleophilic attack is highlighted by a dotted circle. **C.** Helicase activity of the WT and the active site mutants of LcTwinkle. **D.** Fluorescence anisotropy DNA binding of the WT and the active site mutants of LcTwinkle. **E.** DNA binding loops from different subunits of the LcTwinkle-DNA complex take the same configuration. **F.** Structural alignments of ATPase sites across the hexamer suggest that they are superimposable. The five ATPase sites were extracted from the LcTwinkle-DNA complex and subunits at the 5’-side of the DNA in each dimer are aligned. The 5’-side subunit is shown as grey surfaces, and the active site residues on the 3’-side subunits are drawn as cartoons and sticks. **G-H**. Subunit doping experiments of ATPase (**G**) and helicase (**H**) activities with WT and E325Q LcTwinkle.

Careful 3D classification yielded 3 additional structures of LcTwinkle with distinct conformations (Figure 1, F-H and Supplementary Figure 3A). One structure obtained at 7.5 Å resolution contains DNA but exhibits altered CTDs (LcTwinkle-DNA_2_). In LcTwinkle-DNA_2_, the terminal helicase subunit (subunit F) has departed from the subunit E and travelled halfway to the 3’-end of the DNA (Figure 1F and Supplementary Figure 6A). This structure may represent an intermediate state during Twinkle translocation. Although the LcTwinkle sample was prepared in the presence of DNA, two apo LcTwinkle structures were identified, with one in the hexamer form (LcTwinkle_6_) and the other existing as a heptamer (LcTwinkle_7_) (Figure 1, G and H). These structures are similar to what was previously observed with apo Twinkle (33,36–38). The LcTwinkle_6_ stays in a similar non-planar lock-washer configuration as the LcTwinkle-DNA complex (Figure 1G and Supplementary Figure 6B), different from the planar rings formed by gp4 or DnaB in the absence of DNA (50–52). The LcTwinkle_7_ at 8.5 Å resolution contains an enlarged central channel of around 40 Å (Figure 1H), similar as the HsTwinkle heptamer (38). Interestingly, a structural gap is also presented in the LcTwinkle_7_. However, the NTDs are completely disordered in both apo structures, suggesting that the NTDs are highly dynamic without DNA (Figure 1, G and H). It has been reported that in the absence of ssDNA, the NTD conformation is controlled by ATP binding (37). The NTDs are ordered without ATP but become disordered in the presence of ATP (37,38), consistent with our structures in the presence of ATP.

### ATP and DNA binding interfaces of Twinkle

The LcTwinkle helicase contains five intact ATPase sites, each formed by residues from two adjacent subunits (Figure 2A). One Mg^2+^ is associated with each ATP molecule (Supplementary Figure 4E). The Walker A motif from one subunit wraps the triphosphate group of the ATP with its backbone amide groups (Figure 2B). The K301 sidechain on the Walker A motif directly interacts with the γ-phosphate of the ATP. The T302 helps coordinate the Mg^2+^ ion. The D396 on the Walker B motif is around 4 Å to the Mg^2+^ and may coordinate the Mg^2+^ through water-mediated interaction. The mutated Q325 interacts with the γ-phosphate group of the ATP and the Mg^2+^. Two sidechains F501 and R481 sandwich the adenine base of the ATP (Figure 2B and Supplementary Figure 7A). Similar to human Twinkle (34), LcTwinkle is promiscuous and can utilize nucleotides with either adenine or thymine base (Supplementary Figure 7B). In contrast, gp4 prefers dTTP over ATP (53). The R481 on LcTwinkle comes from a different secondary structural element as in gp4 and the F501 is the equivalent of the gp4 Y535 (Supplementary Figure 7A). Mutations of F501A and R481A reduce K_M_ of ATP by ~10-fold (Supplementary Table 2). Surprisingly, F501Y reduces ATP binding as well and makes LcTwinkle favouring the adenine over the thymine base (Supplementary Figure 7B). The K487 and R489 (the arginine finger) from the adjacent subunits contact the γ-phosphate of the ATP. In addition, the H438 sidechain and the Q461 backbone are also in proximity to the γ-phosphate and may stabilize the water for nucleophilic attack during ATP hydrolysis (Figure 2B). Mutations of R489A and H438A significantly reduce the ATPase activity and eliminate DNA binding and helicase activities of LcTwinkle, confirming their critical roles in ATP sensing and translocation (Figure 2, C and D and Supplementary Table 2).

DNA binding in LcTwinkle is mediated by two adjacent loops (LoopD1 and LoopD2 in Figure 2, A and B). The G455 and the A457 backbone amide groups and the S456 sidechain from the LoopD1 and the R440 and the K441 from the LoopD2 interact with the phosphate backbones of DNA (Figure 2B). In addition, the Y412 sidechain contacts the O4’ atom on the sugar ring (Figure 2B). The DNA binding residues in each subunit span four nucleotides distance, with residues on the LoopD1 and the R440 on the LoopD2 interacting with two nucleotides, and the K441 sidechain grabbing two additional nucleotides on the 3’-side of DNA (Figure 2B). DNA bound to the subunit F lacks the interaction with K441 due to the absence of a 5’-end neighbour. Two similar DNA binding loops were found in gp4 helicase (Supplementary Figure 7c) (20). However, the corresponding position of the F454 is an arginine in gp4, which is essential for gp4 DNA binding (54). The LoopD2 in gp4 contains additional positively charged residues and packs on the 3’-side of the coiled ssDNA (Supplementary Figure 7c) (55). Besides directly binding DNA, the positively charged LoopD2 was proposed to facilitate the large-scale domain movement during translocation and prevent DNA from slipping out of gp4 (Supplementary Figure 7D) (26,55). In contrast, the LoopD2 in LcTwinkle is shorter and negatively charged. The DNA is exposed when observed from the C-terminal side of LcTwinkle (Supplementary Figure 7E). Overall, our structure suggested that LcTwinkle is with reduced DNA interactions compared to that of gp4. Direct mutagenesis confirmed the structural model of Twinkle-DNA interaction. Mutations R440A and S456A eliminate DNA binding and helicase activities (Figure 2, C and D). K441A LcTwinkle had low solubility and could not be purified.

Each LcTwinkle helicase subunit is similar within the LcTwinkle hexamer, with an average RMSD of around 0.2 Å (Supplementary Figure 6c). The DNA binding loops are superimposable, except that K441 from subunit A at the DNA 3’-end takes a different conformation (Figure 2E). Moreover, the ATPase sites at the domain interfaces are almost identical (Figure 2F and Supplementary Figure 6D). This is distinct from most other hexameric helicases, where gradual conformational changes of ATPase sites along the helicase hexamer were thought to correspond to sequential ATP hydrolysis and uni-directional translocation (20,22–25). When the planar apo gp4 is used as a reference, one subunit in a LcTwinkle dimer rotates 14 degrees relative to its neighboring subunit (Supplementary Figure 6E). As a comparison, the gp4 subunits rotate 17-23 degrees from the planar conformation (Supplementary Figure 6E). The almost identical configurations of the ATPase sites in the LcTwinkle hexamer suggested that all ATPase sites have a similar chance of hydrolyzing ATP. Moreover, the similar hexameric structures with and without DNA (Supplementary Figure 6B) indicated that DNA binding would not significantly stimulate Twinkle ATPase activity. Notably, gp4 dTTPase activity is stimulated 40-100 folds by DNA, whereas Twinkle ATPase activity is only increased by less than 2 folds upon DNA binding (Supplementary Table 2) (34,49).

To further investigate the mechanism of LcTwinkle ATP hydrolysis and translocation, we performed subunit doping experiments (49,56). When mutated enzymes with defected catalysis were titrated to the WT enzymes, the change in the activity level of the heterooligomer correlates with the subunit cooperativity. If the nucleotide hydrolysis is stochastic within the oligomer, the activity will decrease linearly; on the other hand, when the nucleotide hydrolysis is highly correlated, the activity will change exponentially. Previous subunit doping experiments indicated that the dTTPase activity is highly correlated in gp4 helicase (49), whereas ATP hydrolysis and translocation in archaeal MCM helicase is only moderately correlated (56). A prerequisite for subunit-doping experiments is the efficient formation of the hetero-oligomeric helicase complex. To confirm that different types of LcTwinkle can form proper hetero-oligomers, we constructed and purified LcTwinkle with either N-terminal cyan fluorescent protein (CFP) or yellow fluorescent protein (YFP) (Supplementary Figure 8, A and B). Mixing of the two types of LcTwinkle produced a fluorescence resonance energy transfer (FRET) signal at the wavelength around 536 nm, corresponding to YFP emission, while the CFP peak at ~470 nm was reduced (Supplementary Figure 8C). Moreover, the FRET signal emerged within 2 minutes upon mixing (Supplementary Figure 8D), suggesting efficient and fast hetero-oligomer formation. Similar as in gp4 subunit doping experiments (49), we picked E325Q LcTwinkle, which eliminates ATP hydrolysis but doesn’t affect the ATP or the DNA binding (Figure 2C). When the E325Q LcTwinkle is titrated to the WT LcTwinkle, both the ATPase activity and the helicase activity drop linearly (Figure 2, G and H). The linear decrease suggested a mechanism of stochastic ATP hydrolysis within the LcTwinkle hexamer. Similar linear decreases for the ATPase and helicase activities were observed with H438A and R489A, despite their different roles in ATP hydrolysis and translocation (Supplementary Figure 8). In addition, we tested subunit doping with the DNA binding mutant R440A (Supplementary Figure 8, I and J). The DNA binding affinity is reduced but much slower than the predicted linear decrease when R440A LcTwinkle was titrated. The helicase activity decreases linearly with the increasing amount of the R440A LcTwinkle, similar to those of ATPase site mutants.

### Disease-related Twinkle mutations

Nearly 60 mutations on Twinkle have been implicated in human diseases (Figure 1A and Supplementary Figure 1C) (11). Excepting four mutations on the extreme N- and C-terminus, all other disease-related mutations on human Twinkle can be mapped onto the LcTwinkle structure, with most of them conserved (Figure 3 and Supplementary Figure 1C).

**Figure 3.**
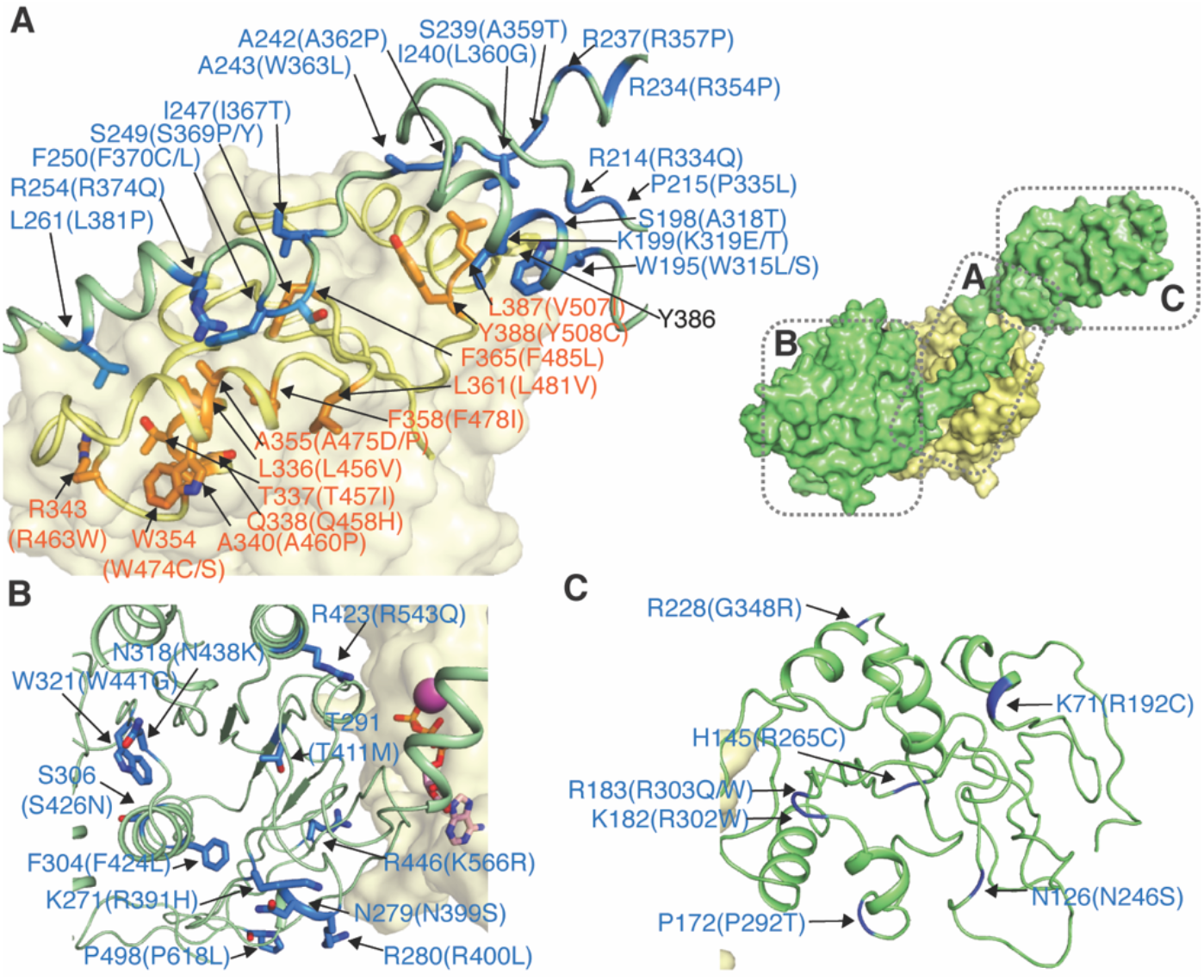
Disease-related mutations on LcTwinkle. **A.** Disease-related mutations on the N-C interface and the N-C linker. Two adjacent subunits C (green) and D (yellow) are selected to illustrate the positions of the disease-related mutations. The mutations on the subunit C are highlighted in blue and the mutations on the subunit D are highlighted in orange. The corresponding human disease-related mutations are indicated in the bracket. The locations of each panel relative to the overall structure were indicated on the small diagram on the up-right corner in (**A**). **B.** Disease-related mutations on the CTD. **C.** Disease-related mutations on the NTD.

The N-C linker and the NTD form extensive interactions with the CTD of its 3’-side subunit (Figure 3A). The domain-swapped interface is a hotspot for disease-related mutations (Figure 3A). The NTD directly contacts the CTD with an α-helix formed by residues 195-204 (Figure 3A and Supplementary Figure 9A). Although the overall resolution is low for the NTDs, the helix is well ordered, and densities for the W195 and the K199 sidechains are visible (Supplementary Figure 4A, 9A). The W195 is in proximity to the Y386 sidechain, and the K199 interacts with the backbone oxygen of the L387 (Supplementary Figure 9A). Residues near the interface, including W195, S198, K199, R214, P215 on the NTD, L387, and Y388 on the CTD, have been found in human patients (Figure 3A). Although Y386 mutation has not been correlated with human disease, Y386A reduces the helicase activity similar to the disease-related mutations K199E, W195L, and Y388C (Supplementary Figure 9B). The N-C interface may stabilize Twinkle hexamer during translocation (37). WT LcTwinkle forms hexamers and heptamers on the negative staining EM grid (Supplementary Figure 9, C and E). In contrast, Y386A, K199E, W195L, and Y388C LcTwinkle samples contain significantly reduced oligomers, with the significant oligomer form of these mutants being heptamers or larger (Supplementary Figure 9, C-J), similar as observed for HsTwinkle (37,38). In contrast, the NTD are located at different places relative to the CTD and the N-C interface is not preserved in the apo HsTwinkle structure (Supplementary Figure 9K), possibly due to the W315L mutation (corresponding to LcTwinkle W195L) on the N-C interface (38).

The N-C linker near the NTD is sandwiched by the NTD and the CTD (Figure 3A). Six mutations on the NTD-linker junction are related to human diseases. The C-terminal end of the N-C linker forms a helix and stacks in a hydrophobic groove on the CTD (Figure 3A). L261, F250, and I247 interact with the CTD through hydrophobic interactions, while R254 forms salt links with D352 (Figure 3A). Mutations of L261, R254, F250, S249, and I247 on the N-C linker and R343, A340, A355, L336, T337, Q338, W354, F358, L361, and F365 on the CTD cause human diseases. The linker in the HsTwinkle structure interacts with the CTD through the same interface, although the linker helix is lifted ~1 Å and the interface is reduced (Supplementary Figure 9J). The N-C linker may play a similar role as the NTDs in stabilizing Twinkle hexamer, as suggested by previous biochemical and structural data (37).

Besides mutations at the domain-swapped interface, eleven mutations are located on the CTD (Figure 3B). Residues W321, N318, S306, and F304 are at the core of the CTD and may be important for CTD folding and stability. R423, T291, R446, N279, and R280 are near the subunit-subunit interface on the CTD. These residues may contribute to the CTD subunit-subunit interactions and ATP hydrolysis. In addition, seven mutations are mapped to the surface of the NTD, and most of them are charged residues (Figure 3C). Of notice, A243 (human disease W363L), R228 (human disease G348R), and R446 (human disease K566R) are different from the corresponding human residues. The differences in A243 and R228 are possibly due to changes in their surrounding residues (Supplementary Figure 9, L and M). R446 is part of the LoopD2 but does not directly contact DNA (Supplementary Figure 7C). How human K566R mutation would affect Twinkle function is unclear.

### HS-AFM imaging of LcTwinkle DNA binding dynamics

Our recent AFM imaging in liquids showed that human Twinkle subunits could self-assemble into hexamers and higher-order complexes and switch between open and closed configurations (57). However, due to the low temporal resolution of the conventional AFM imaging, we could not elucidate the real-time conformational changes of Twinkle during DNA binding. To overcome this technical barrier, we used HS-AFM imaging in liquids to simultaneously convene structural and dynamic information of LcTwinkle. A linear DNA (2030 bps) (57) and the WT LcTwinkle were deposited onto a 1-(3-Aminopropyl) silatrane (APS)-treated mica (APS-mica) surface (58) in the presence of ATP. The AFM images were recorded at ~0.8-2 frames/s to visualize the dynamics of the LcTwinkle. Under this imaging condition, both DNA and LcTwinkle were mobile on the APS-mica surface.

In the absence of a proximal DNA molecule (LcTwinkle-DNA distance > 200 nm), the LcTwinkle exhibited limited conformational changes manifested as opening and closing of gaps between subunits (Supplementary Video 1; panels I and II in Figure 4A and Supplementary Video 2; N=17 Twinkle moledules), similar to human Twinkle (57). Unexpectedly, when DNA molecules were present in the vicinity of LcTwinkle (LcTwinkle-DNA distance < 20 nm), configurations of the LcTwinkle changed dramatically (Figure 4A and Supplementary Video 2). A domain from the LcTwinkle oligomer proximal to the DNA protruded out to capture the nearby DNA (Figure 4, A-D and Supplementary Videos 2-6). Domain protrusions were evident in all LcTwinkle molecules proximal to DNA (mean LcTwinkle-DNA distance 10.7 ± 3.7 nm, N=27, Figure 4E). The average length of the protrusion was 4.9 ± 1.8 nm (Figure 4E, N=27). When DNA moved around or multiple DNA segments were nearby, we observed frequent domain protrusions from multiple subunits of the LcTwinkle (Supplementary Figure 10 and Supplementary Video 4). In addition, during the DNA search, some Twinkle rings were stretched to an almost linear form (Figure 4C and Supplementary Video 5). HS-AFM imaging also revealed the capture and the release of DNA at the central channel of the LcTwinkle (Figure 4D and Supplementary Videos 6,7; N=7 events). The DNA binding through domain protrusion appeared to proceed with the loading of DNA into the central channel (Figure 4D, Panels I-IV). It is worth noting that the ring-opening of the LcTwinkle allowed the entering or release of DNA from its central channel (Figure 4D, Panels III and V). In addition, after DNA left the central channel, LcTwinkle transiently interacted with the protruded domain again (Figure 4D, Panels VII and VIII).

**Figure 4.**
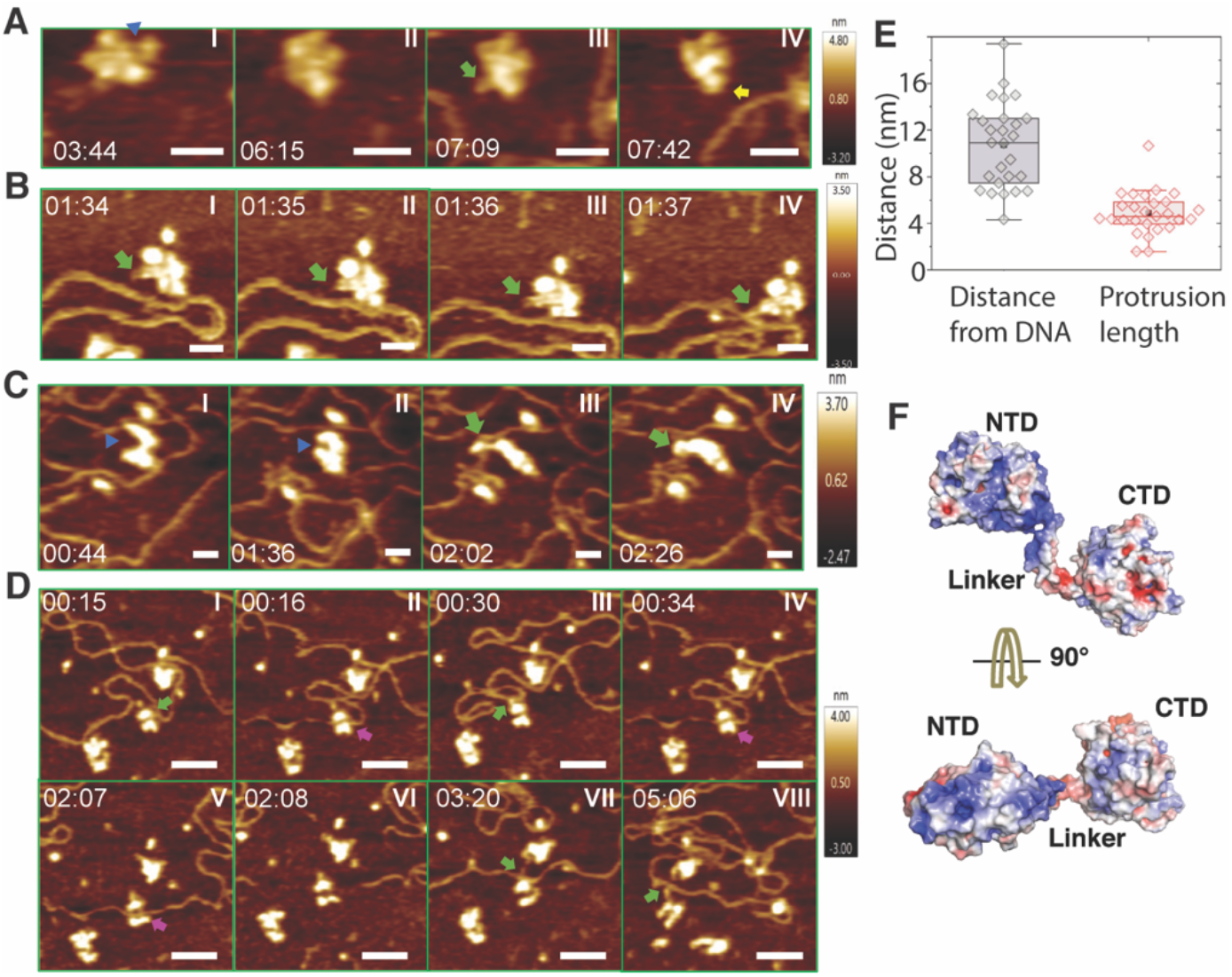
Real-time HS-AFM imaging of LcTwinkle shows DNA capture by N-protrusion and at the central channel. **A.** LcTwinkle switches between open (blue triangle) and closed conformations in the absence of DNA (Panels I and II). Nearby DNA induces LcTwinkle N-protrusion (Panels III and IV). The N-protrusion is highlighted by green (N-protrusion on X- and Y-plane) and yellow (N-protrusion characterized by an increased height - brighter Z scale) arrows. Also, see Supplementary Video 2. XY Scale bar = 20 nm. **B-C**. LcTwinkle conformers showing N-protrusions that capture DNA in the vicinity (distance to DNA less than 20 nm). Also, see Supplementary Videos 3 and 5. The open gap in LcTwinkle is indicated as blue triangles in Panels I and II in panel C. XY Scale bar = 20 nm. **D**. DNA capture and release revealed by HS-AFM imaging. The green and purple arrows highlight DNA binding at the N-protrusion and the CTD central channel, respectively. Also, see Supplementary Video 6. XY Scale bar = 50 nm. **E**. Box plots showing the distance distribution between DNA and the protruded subunit on LcTwinkle (10.7 nm ± 3.7 nm, N = 27) and the protruded length of the LcTwinkle conformers (4.9 nm ± 1.8 nm, N = 27). The reported mean and standard deviation are from HS-AFM images collected from three independent experiments. **F**. Two views of the electrostatic surfaces of a LcTwinkle subunit. The negatively charged surfaces are colored blue, and the positively charged surfaces are colored red.

## DISCUSSION

Most replicative helicases contain NTDs in addition to the C-terminal helicase domains (Supplementary Figure 5) (20–23,25). The gp4 NTD encodes a primase domain, whereas the NTDs in bacterial, archaeal, and eukaryotic replicative helicases physically interact with the primases (50,59). The NTDs are always on the 5’-side of the DNA relative to the helicase domains to assist primer synthesis (Supplementary Figure 5). Like gp4, Twinkle helicases from plants and lower eukaryotes can catalyze primer synthesis (60,61). However, the primase active site residues are mutated in vertebrate Twinkle (11). Furthermore, the NTD rotates 90° and flips 180° away from the unwound DNA (Supplementary Figure 5I). The unique NTD arrangement is consistent with the lack of Okazaki fragment synthesis in animal mitochondria (3,4). Moreover, the NTD conformations are likely under the dual control of both DNA and ATP. Without DNA and ATP, the NTD is observed to attach to the CTD of the same subunit in HsTwinkle (38). Without DNA, NTD becomes disordered in the presence of ATP (37). Simultaneous binding of ATP and DNA stabilizes NTD conformation.

Although without enzymatic activities, the Twinkle NTDs likely play essential roles in Twinkle oligomerization, additional DNA binding, and protein-protein interactions, similar to the NTDs in gp4 and DnaB (50,62). Our LcTwinkle structure shows that the NTD and the N-C linker directly interact with the CTD in its neighbouring subunit in the DNA bound state (Figure 1D, 3A). The interaction stabilizes Twinkle hexamers around DNA. Mutations at the interface alter oligomerization of Twinkle and reduce helicase unwinding (Supplementary Figure 9) (37). The domain-swapped interface is a hotspot for disease-related mutations (Figure 3A). In addition, the NTD may facilitate Twinkle DNA capture and loading (as discussed below). Moreover, strand annealing and exchange activities have been reported for human Twinkle (34), and these activities require more than one DNA binding interface. The six NTDs may capture multiple ssDNA molecules to assist their pairing. In addition, Twinkle is reported to interact with multiple factors in DNA replication and repair (6,15–17). Any proteins operating on the lagging strand DNA may contact Twinkle through the NTD. Seven disease-related mutations are mapped onto the NTD. They are far away from the CTD or the NTD-CTD interface, thus not likely to be directly involved in Twinkle unwinding. These mutations may affect Twinkle protein or DNA interactions.

Loading of replicative helicases onto the genomic DNA is a prerequisite and often a critical regulatory step in DNA replication (28). In bacteria, archaeal, and eukaryotic nucleus, an origin recognition complex recognizes the replication origin and recruits the helicase. In addition, a helicase loader helps open the helicase ring for loading (28). Neither origin recognition complex nor helicase loader has been identified in mitochondria. Twinkle itself must search and capture the DNA substrate in 3-dimensional space. Remarkably, HS-AFM imaging in liquids uncovered a novel proximal DNA-induced conformational change of Twinkle. Using HS-AFM, we observed that when LcTwinkle is close to DNA (LcTwinkle-DNA distance < 20 nm), a domain could move freely in solution and protrude approximately 5 nm in length, away from the major portion of LcTwnk oligomers to search and capture nearby DNA (Figure 4). We propose that the mobile domain is the NTD. The NTDs do not interact with each other (Figure. 1d), while the CTDs can oligomerize in the presence of ATP, which was included in our AFM imaging buffer. Our apo LcTwinkle and previous apo HsTwinkle structures (38) confirmed the dynamic nature of the NTDs (Fig. 1, G and H). The N-C linker spans over 30 residues and can account for the long-distance protrusion when it becomes fully extended. Therefore, we termed the domain protrusion observed in HS-AFM as “N-protrusion” (Figure 4 and Supplementary Videos 2-6). The N-protrusion is likely guided by electrostatic interactions between the NTDs and DNA. There are several positively charged patches on the NTD that may potentially interact with the negatively charged DNA electrostatically over long distances (Figure 4F). It is worth noting that the Debye screening length around dsDNA is ~1 nm at the ionic strength used in HS-AFM imaging (63). Since both proteins and DNA were mobile during AFM imaging, the precise distance between LcTwinkle and DNA that activates N-protrusion could be significantly shorter than what we measure based on individual AFM images. The length of N-protrusion could also be augmented by mobile DNA attached to the NTD.

The extraordinary conformational change of the N-protrusion suggests a plausible mechanism for Twinkle DNA loading in the absence of a helicase loader (Fig. 5). Individual NTDs from an apo Twinkle molecule either sit on its own CTD without domain swap (38) or extend out to search for nearby DNA (Figure 4 and Supplementary Videos 2-6). After DNA capture, the protruded NTD helps bring the DNA into its central channel for loading (Figure 4D and Supplementary Video 6). In addition, the Twinkle hexameric ring connected by the N-C linkers must open to allow DNA entrance. Apo LcTwinkle is likely prone to ring opening due to the lack of domain-swapped N-C interactions. HS-AFM revealed that LcTwinkle could switch between open and closed-ring conformations in the absence of DNA (Figure 4A, panels I and II, and Supplementary Video 2). Our HS-AFM data also confirmed that Twinkle could capture DNA at its central channel through ring-opening (Figure 4D and Supplementary Videos 6,7) without subunit association or dissociation. Taken together, the cryo-EM and HS-AFM data suggest two separate DNA binding regions on Twinkle, the NTD for initial DNA capture and the helicase central channel for DNA unwinding. Our model of Twinkle loading (Fig. 5) is also consistent with a model proposed for gp4 helicase, which involves the initial binding of the DNA to the NTD outside of the helicase ring, a conformational transition, followed by the migration of the DNA into the central channel, and ring closure (30). Although not directly observed, we cannot exclude alternative possibilities of LcTwinkle loading by assembling of monomers or dissociation of a subunit from a heptamer.

**Figure 5.**
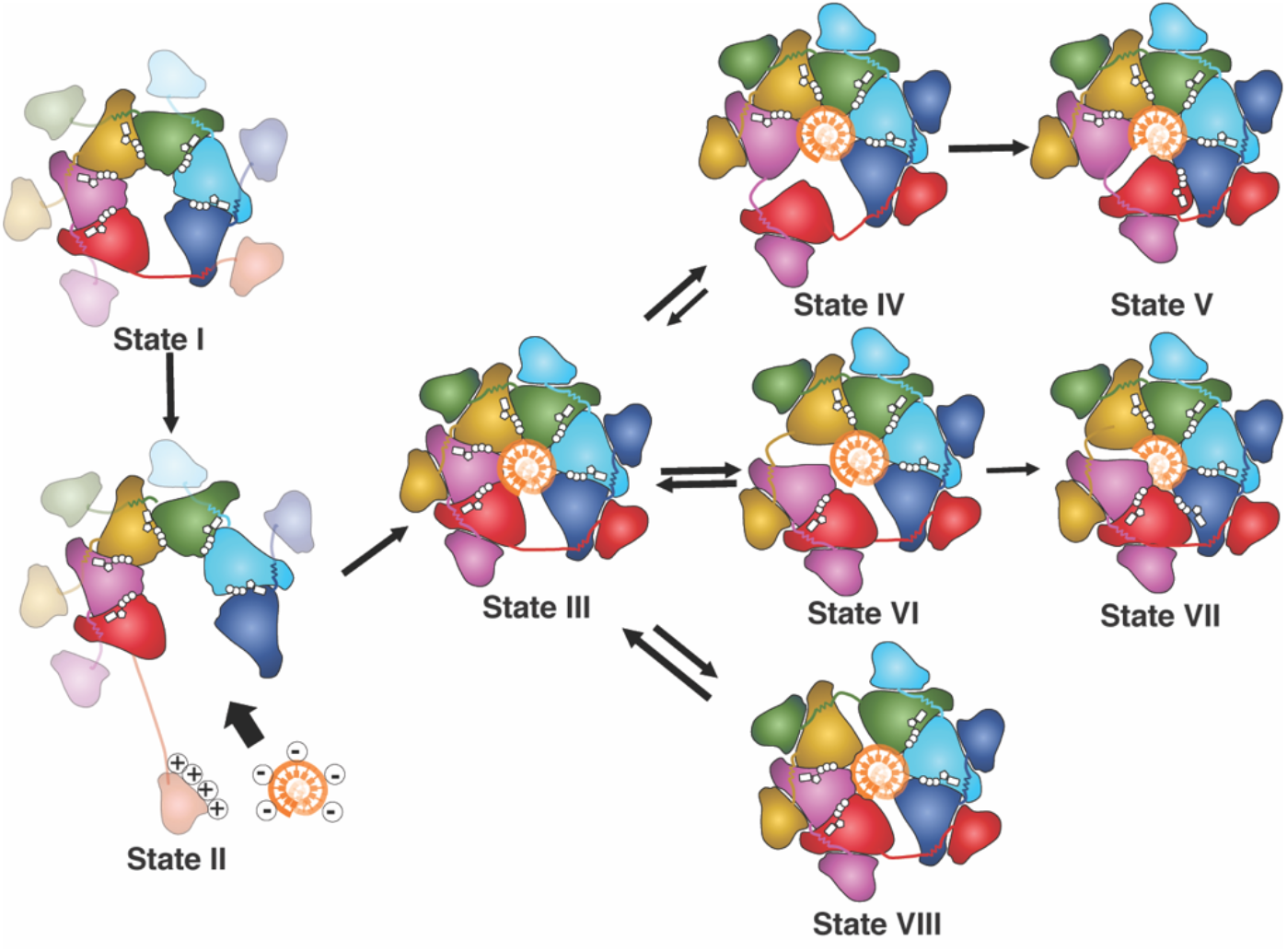
Model of LcTwinkle loading and translocation. Twinkle in the absence of DNA is prone to ring-opening (State I). When DNA is nearby, the positively charged NTD captures the negatively charged DNA over long-distance and the CTD hexamer opens further for DNA loading (State II). After loading, Twinkle forms a lock-washer-shaped hexamer with the NTDs attaching to the side of the CTDs and the DNA binding to the CTD central channel (State III). ATP hydrolysis is stochastic in Twinkle. When ATP is hydrolyzed at the subunit interfaces near the DNA 5’-end (State IV and State VI), one or two subunits dissociate from its neighbor and travels to the other end of the DNA (State V and State VII). However, when ATP is hydrolyzed in the middle or close to the DNA 3’-end (State VIII), the translocation is unfavored, leading to a futile cycle of ATP hydrolysis.

The ATP binding and hydrolysis empower helicase translocation on DNA. While several models of ATP hydrolysis have been proposed in hexameric ATPases, the sequential ATP hydrolysis model is widely accepted (64). In contrast, our structural and biochemical analyses suggest that the ATP hydrolysis is stochastic in the LcTwinkle. The random ATP hydrolysis was also reported for the hexameric ClpX peptide translocase, where the loss of one or more ATPase sites only reduces but does not eliminate translocation (65). Similarly, ATPase sites in the hetero-hexameric CMG helicase are not all required for its translocation (25). Helicase translocation is associated with frequent futile cycles, slippering, and backtracking (53). Possibly, ATP hydrolysis near the 5’-end of DNA will lead to translocation of one to two subunits, whereas ATP hydrolysis in the middle or close to the 3’-end may result in futile ATPase cycles (Fig. 5). The *en bloc* movement of multiple subunits also explains how inactive subunits can be tolerated in Twinkle, CMG, and ClpX. Twinkle can move on DNA unidirectionally in the 5’-to-3’ direction (31). Our LcTwinkle-DNA structure suggested that subunit F on the 5’-end of DNA is with reduced DNA binding, as one of the key DNA binding residues K441 from a neighbour subunit is missing at the 5’-end of DNA. It is possible that the subtle difference in DNA binding renders subunit F most mobile. On the other hand, the directional movement could be affected by the NTDs. SF4 G40P helicase is with minimal NTD and can translocate in both directions on ssDNA (66). In addition, a comparison of DNA binding loops in gp4 and LcTwinkle also suggested that DNA may easily slide out from the Twinkle DNA binding channel (Supplementary Figure 7E). Thus, our structural and biochemical suggest that Twinkle is inefficient in unwinding DNA due to the stochastic ATP hydrolysis and reduced DNA binding. Indeed, Twinkle ATP hydrolysis is 20-100 folds slower than that of gp4 or DnaB (34). It is possible that Twinkle only stays in a partially active form and additional binding partners or post-translational modifications fully activate Twinkle.

In summary, our biochemical, biophysical and structural data illustrate unique structural and dynamic features of the mitochondrial replicative helicase Twinkle. Our data highlight the important role of the enzymatically inactive NTD in Twinkle loading, unwinding, and the NTD-CTD interface represents a hotspot for human disease related mutations.

## Supporting information

Combined supplementary material

## DATA AVAILABILITY

All original data and materials will be available upon request.

## ACCESSION NUMBERS

The three-dimensional cryo-EM density maps for LcTwinkle complexes have been deposited in the EM Database under the accession code EMDB: EMD-XXXX, EMD-XXXX, EMD-XXXX and EMD-XXXX, and the coordinates for the structure have been deposited in Protein Data Bank under accession code PDB XXXX.

## SUPPLEMENTARY DATA

Supplementary Data are available at NAR online.

## ACKNOWLEDGEMENTS

We thank Wenhua Guo from Rice University Shared Equipment Authority, Isaac Forester from cryo-EM core facility at Baylor College of Medicine (supported by CPRIT RP190602), Venkata Mallmapalli, and Guizhen Fan from cryo-EM core facility at the University of Texas McGovern Medical School (supported by CPRIT RP190602) for their help in cryo-EM sample preparation, screening and data collection. We thank Ryan Fuierer and Keith Jones at Asylum Research for access to VRS AFM. We thank Dr. Wei Yang from NIH for her support in the initial clone of the LcTwinkle gene.

## REFERENCES

1. Robberson, D.L., Kasamatsu, H. and Vinograd, J. (1972) Replication of mitochondrial DNA. Circular replicative intermediates in mouse L cells. Proc Natl Acad Sci U S A, 69, 737–741.

2. Gustafsson, C.M., Falkenberg, M. and Larsson, N.G. (2016) Maintenance and Expression of Mammalian Mitochondrial DNA. Annu Rev Biochem, 85, 133–160.

3. Yasukawa, T., Reyes, A., Cluett, T.J., Yang, M.Y., Bowmaker, M., Jacobs, H.T. and Holt, I.J. (2006) Replication of vertebrate mitochondrial DNA entails transient ribonucleotide incorporation throughout the lagging strand. EMBO J, 25, 5358–5371.

4. Miralles Fuste, J., Shi, Y., Wanrooij, S., Zhu, X., Jemt, E., Persson, O., Sabouri, N., Gustafsson, C.M. and Falkenberg, M. (2014) In vivo occupancy of mitochondrial single-stranded DNA binding protein supports the strand displacement mode of DNA replication. PLoS Genet, 10, e1004832.

5. Falkenberg, M. (2018) Mitochondrial DNA replication in mammalian cells: overview of the pathway. Essays Biochem, 62, 287–296.

6. Korhonen, J.A., Pham, X.H., Pellegrini, M. and Falkenberg, M. (2004) Reconstitution of a minimal mtDNA replisome in vitro. EMBO J, 23, 2423–2429.

7. Wanrooij, S., Fuste, J.M., Farge, G., Shi, Y., Gustafsson, C.M. and Falkenberg, M. (2008) Human mitochondrial RNA polymerase primes lagging-strand DNA synthesis in vitro. Proc Natl Acad Sci U S A, 105, 11122–11127.

8. Fuste, J.M., Wanrooij, S., Jemt, E., Granycome, C.E., Cluett, T.J., Shi, Y., Atanassova, N., Holt, I.J., Gustafsson, C.M. and Falkenberg, M. (2010) Mitochondrial RNA polymerase is needed for activation of the origin of light-strand DNA replication. Mol Cell, 37, 67–78.

9. DeBalsi, K.L., Hoff, K.E. and Copeland, W.C. (2017) Role of the mitochondrial DNA replication machinery in mitochondrial DNA mutagenesis, aging and age-related diseases. Ageing Res Rev, 33, 89–104.

10. Spelbrink, J.N., Li, F.Y., Tiranti, V., Nikali, K., Yuan, Q.P., Tariq, M., Wanrooij, S., Garrido, N., Comi, G., Morandi, L. et al. (2001) Human mitochondrial DNA deletions associated with mutations in the gene encoding Twinkle, a phage T7 gene 4-like protein localized in mitochondria. Nat Genet, 28, 223–231.

11. Peter, B. and Falkenberg, M. (2020) TWINKLE and Other Human Mitochondrial DNA Helicases: Structure, Function and Disease. Genes (Basel), 11.

12. Tyynismaa, H., Sembongi, H., Bokori-Brown, M., Granycome, C., Ashley, N., Poulton, J., Jalanko, A., Spelbrink, J.N., Holt, I.J. and Suomalainen, A. (2004) Twinkle helicase is essential for mtDNA maintenance and regulates mtDNA copy number. Hum Mol Genet, 13, 3219–3227.

13. Baloh, R.H., Salavaggione, E., Milbrandt, J. and Pestronk, A. (2007) Familial parkinsonism and ophthalmoplegia from a mutation in the mitochondrial DNA helicase twinkle. Arch Neurol, 64, 998–1000.

14. Pohjoismaki, J.L., Williams, S.L., Boettger, T., Goffart, S., Kim, J., Suomalainen, A., Moraes, C.T. and Braun, T. (2013) Overexpression of Twinkle-helicase protects cardiomyocytes from genotoxic stress caused by reactive oxygen species. Proc Natl Acad Sci U S A, 110, 19408–19413.

15. Sykora, P., Kanno, S., Akbari, M., Kulikowicz, T., Baptiste, B.A., Leandro, G.S., Lu, H., Tian, J., May, A., Becker, K.A. et al. (2017) DNA Polymerase Beta Participates in Mitochondrial DNA Repair. Mol Cell Biol, 37.

16. Peeva, V., Blei, D., Trombly, G., Corsi, S., Szukszto, M.J., Rebelo-Guiomar, P., Gammage, P.A., Kudin, A.P., Becker, C., Altmuller, J. et al. (2018) Linear mitochondrial DNA is rapidly degraded by components of the replication machinery. Nat Commun, 9, 1727.

17. Stojkovic, G., Makarova, A.V., Wanrooij, P.H., Forslund, J., Burgers, P.M. and Wanrooij, S. (2016) Oxidative DNA damage stalls the human mitochondrial replisome. Sci Rep, 6, 28942.

18. Gao, Y. and Yang, W. (2020) Different mechanisms for translocation by monomeric and hexameric helicases. Curr Opin Struct Biol, 61, 25–32.

19. Singleton, M.R., Dillingham, M.S. and Wigley, D.B. (2007) Structure and mechanism of helicases and nucleic acid translocases. Annu Rev Biochem, 76, 23–50.

20. Gao, Y., Cui, Y., Fox, T., Lin, S., Wang, H., de Val, N., Zhou, Z.H. and Yang, W. (2019) Structures and operating principles of the replisome. Science, 363.

21. Itsathitphaisarn, O., Wing, R.A., Eliason, W.K., Wang, J. and Steitz, T.A. (2012) The hexameric helicase DnaB adopts a nonplanar conformation during translocation. Cell, 151, 267–277.

22. Enemark, E.J. and Joshua-Tor, L. (2006) Mechanism of DNA translocation in a replicative hexameric helicase. Nature, 442, 270–275.

23. Meagher, M., Epling, L.B. and Enemark, E.J. (2019) DNA translocation mechanism of the MCM complex and implications for replication initiation. Nat Commun, 10, 3117.

24. Thomsen, N.D. and Berger, J.M. (2009) Running in reverse: the structural basis for translocation polarity in hexameric helicases. Cell, 139, 523–534.

25. Eickhoff, P., Kose, H.B., Martino, F., Petojevic, T., Abid Ali, F., Locke, J., Tamberg, N., Nans, A., Berger, J.M., Botchan, M.R. et al. (2019) Molecular Basis for ATP-Hydrolysis-Driven DNA Translocation by the CMG Helicase of the Eukaryotic Replisome. Cell Rep, 28, 2673-2688 e2678.

26. Jin, S., Bueno, C., Lu, W., Wang, Q., Chen, M., Chen, X., Wolynes, P.G. and Gao, Y. (2022) Computationally exploring the mechanism of bacteriophage T7 gp4 helicase translocating along ssDNA. Proc Natl Acad Sci U S A, 119, e2202239119.

27. Puchades, C., Sandate, C.R. and Lander, G.C. (2020) The molecular principles governing the activity and functional diversity of AAA+ proteins. Nat Rev Mol Cell Biol, 21, 43–58.

28. Bleichert, F., Botchan, M.R. and Berger, J.M. (2017) Mechanisms for initiating cellular DNA replication. Science, 355.

29. Jemt, E., Farge, G., Backstrom, S., Holmlund, T., Gustafsson, C.M. and Falkenberg, M. (2011) The mitochondrial DNA helicase TWINKLE can assemble on a closed circular template and support initiation of DNA synthesis. Nucleic Acids Res, 39, 9238–9249.

30. Ahnert, P., Picha, K.M. and Patel, S.S. (2000) A ring-opening mechanism for DNA binding in the central channel of the T7 helicase-primase protein. EMBO J, 19, 3418–3427.

31. Korhonen, J.A., Gaspari, M. and Falkenberg, M. (2003) TWINKLE Has 5’-> 3’ DNA helicase activity and is specifically stimulated by mitochondrial single-stranded DNA-binding protein. J Biol Chem, 278, 48627–48632.

32. Shutt, T.E. and Gray, M.W. (2006) Bacteriophage origins of mitochondrial replication and transcription proteins. Trends Genet, 22, 90–95.

33. Fernandez-Millan, P., Lazaro, M., Cansiz-Arda, S., Gerhold, J.M., Rajala, N., Schmitz, C.A., Silva-Espina, C., Gil, D., Bernado, P., Valle, M. et al. (2015) The hexameric structure of the human mitochondrial replicative helicase Twinkle. Nucleic Acids Res, 43, 4284–4295.

34. Sen, D., Nandakumar, D., Tang, G.Q. and Patel, S.S. (2012) Human mitochondrial DNA helicase TWINKLE is both an unwinding and annealing helicase. J Biol Chem, 287, 14545–14556.

35. Khan, I., Crouch, J.D., Bharti, S.K., Sommers, J.A., Carney, S.M., Yakubovskaya, E., Garcia-Diaz, M., Trakselis, M.A. and Brosh, R.M., Jr. (2016) Biochemical Characterization of the Human Mitochondrial Replicative Twinkle Helicase: SUBSTRATE SPECIFICITY, DNA BRANCH MIGRATION, AND ABILITY TO OVERCOME BLOCKADES TO DNA UNWINDING. J Biol Chem, 291, 14324–14339.

36. Ziebarth, T.D., Gonzalez-Soltero, R., Makowska-Grzyska, M.M., Nunez-Ramirez, R., Carazo, J.M. and Kaguni, L.S. (2010) Dynamic effects of cofactors and DNA on the oligomeric state of human mitochondrial DNA helicase. J Biol Chem, 285, 14639–14647.

37. Peter, B., Farge, G., Pardo-Hernandez, C., Tangefjord, S. and Falkenberg, M. (2019) Structural basis for adPEO-causing mutations in the mitochondrial TWINKLE helicase. Hum Mol Genet, 28, 1090–1099.

38. Riccio, A.A., Bouvette, J., Perera, L., Longley, M.J., Krahn, J.M., Williams, J.G., Dutcher, R., Borgnia, M.J. and Copeland, W.C. (2022) Structural insight and characterization of human Twinkle helicase in mitochondrial disease. Proc Natl Acad Sci U S A, 119, e2207459119.

39. Edgar, R.C. (2004) MUSCLE: multiple sequence alignment with high accuracy and high throughput. Nucleic Acids Res, 32, 1792–1797.

40. Zivanov, J., Nakane, T., Forsberg, B.O., Kimanius, D., Hagen, W.J., Lindahl, E. and Scheres, S.H. (2018) New tools for automated high-resolution cryo-EM structure determination in RELION-3. Elife, 7.

41. Zheng, S.Q., Palovcak, E., Armache, J.P., Verba, K.A., Cheng, Y. and Agard, D.A. (2017) MotionCor2: anisotropic correction of beam-induced motion for improved cryo-electron microscopy. Nat Methods, 14, 331–332.

42. Zhang, K. (2016) Gctf: Real-time CTF determination and correction. J Struct Biol, 193, 1–12.

43. Punjani, A., Rubinstein, J.L., Fleet, D.J. and Brubaker, M.A. (2017) cryoSPARC: algorithms for rapid unsupervised cryo-EM structure determination. Nat Methods, 14, 290–296.

44. Pettersen, E.F., Goddard, T.D., Huang, C.C., Couch, G.S., Greenblatt, D.M., Meng, E.C. and Ferrin, T.E. (2004) UCSF Chimera--a visualization system for exploratory research and analysis. J Comput Chem, 25, 1605–1612.

45. Emsley, P., Lohkamp, B., Scott, W.G. and Cowtan, K. (2010) Features and development of Coot. Acta Crystallogr D Biol Crystallogr, 66, 486–501.

46. Afonine, P.V., Poon, B.K., Read, R.J., Sobolev, O.V., Terwilliger, T.C., Urzhumtsev, A. and Adams, P.D. (2018) Real-space refinement in PHENIX for cryo-EM and crystallography. Acta Crystallogr D Struct Biol, 74, 531–544.

47. Countryman, P., Fan, Y., Gorthi, A., Pan, H., Strickland, J., Kaur, P., Wang, X., Lin, J., Lei, X., White, C. et al. (2018) Cohesin SA2 is a sequence-independent DNA-binding protein that recognizes DNA replication and repair intermediates. J Biol Chem, 293, 1054–1069.

48. Ziebarth, T.D., Farr, C.L. and Kaguni, L.S. (2007) Modular architecture of the hexameric human mitochondrial DNA helicase. J Mol Biol, 367, 1382–1391.

49. Crampton, D.J., Mukherjee, S. and Richardson, C.C. (2006) DNA-induced switch from independent to sequential dTTP hydrolysis in the bacteriophage T7 DNA helicase. Mol Cell, 21, 165–174.

50. Bailey, S., Eliason, W.K. and Steitz, T.A. (2007) Structure of hexameric DnaB helicase and its complex with a domain of DnaG primase. Science, 318, 459–463.

51. Singleton, M.R., Sawaya, M.R., Ellenberger, T. and Wigley, D.B. (2000) Crystal structure of T7 gene 4 ring helicase indicates a mechanism for sequential hydrolysis of nucleotides. Cell, 101, 589–600.

52. Toth, E.A., Li, Y., Sawaya, M.R., Cheng, Y. and Ellenberger, T. (2003) The crystal structure of the bifunctional primase-helicase of bacteriophage T7. Mol Cell, 12, 1113–1123.

53. Sun, B., Johnson, D.S., Patel, G., Smith, B.Y., Pandey, M., Patel, S.S. and Wang, M.D. (2011) ATP-induced helicase slippage reveals highly coordinated subunits. Nature, 478, 132–135.

54. Washington, M.T., Rosenberg, A.H., Griffin, K., Studier, F.W. and Patel, S.S. (1996) Biochemical analysis of mutant T7 primase/helicase proteins defective in DNA binding, nucleotide hydrolysis, and the coupling of hydrolysis with DNA unwinding. J Biol Chem, 271, 26825–26834.

55. Satapathy, A.K., Kochaniak, A.B., Mukherjee, S., Crampton, D.J., van Oijen, A. and Richardson, C.C. (2010) Residues in the central beta-hairpin of the DNA helicase of bacteriophage T7 are important in DNA unwinding. Proc Natl Acad Sci U S A, 107, 6782–6787.

56. Moreau, M.J., McGeoch, A.T., Lowe, A.R., Itzhaki, L.S. and Bell, S.D. (2007) ATPase site architecture and helicase mechanism of an archaeal MCM. Mol Cell, 28, 304–314.

57. Kaur, P., Longley, M.J., Pan, H., Wang, W., Countryman, P., Wang, H. and Copeland, W.C. (2020) Single-molecule level structural dynamics of DNA unwinding by human mitochondrial Twinkle helicase. Journal of Biological Chemistry, 295, 5564–5576.

58. Shlyakhtenko, L.S., Gall, A.A. and Lyubchenko, Y.L. (2013) Mica functionalization for imaging of DNA and protein-DNA complexes with atomic force microscopy. Methods in molecular biology, 931, 295–312.

59. Sun, J., Shi, Y., Georgescu, R.E., Yuan, Z., Chait, B.T., Li, H. and O’Donnell, M.E. (2015) The architecture of a eukaryotic replisome. Nat Struct Mol Biol, 22, 976–982.

60. Harman, A. and Barth, C. (2018) The Dictyostelium discoideum homologue of Twinkle, Twm1, is a mitochondrial DNA helicase, an active primase and promotes mitochondrial DNA replication. BMC Mol Biol, 19, 12.

61. Diray-Arce, J., Liu, B., Cupp, J.D., Hunt, T. and Nielsen, B.L. (2013) The Arabidopsis At1g30680 gene encodes a homologue to the phage T7 gp4 protein that has both DNA primase and DNA helicase activities. BMC Plant Biol, 13, 36.

62. Yang, W., Seidman, M.M., Rupp, W.D. and Gao, Y. (2019) Replisome structure suggests mechanism for continuous fork progression and post-replication repair. DNA Repair (Amst), 81, 102658.

63. Sorgenfrei, S., Chiu, C.Y., Johnston, M., Nuckolls, C. and Shepard, K.L. (2011) Debye screening in single-molecule carbon nanotube field-effect sensors. Nano Lett, 11, 3739–3743.

64. Lyubimov, A.Y., Strycharska, M. and Berger, J.M. (2011) The nuts and bolts of ring-translocase structure and mechanism. Curr Opin Struct Biol, 21, 240–248.

65. Martin, A., Baker, T.A. and Sauer, R.T. (2005) Rebuilt AAA + motors reveal operating principles for ATP-fuelled machines. Nature, 437, 1115–1120.

66. Mesa, P., Alonso, J.C. and Ayora, S. (2006) Bacillus subtilis bacteriophage SPP1 G40P helicase lacking the n-terminal domain unwinds DNA bidirectionally. J Mol Biol, 357, 1077–1088.

