## Supplementary material for "Structural and Dynamic Basis of DNA Capture and Translocation by Mitochondrial Twinkle Helicase": Combined supplementary material

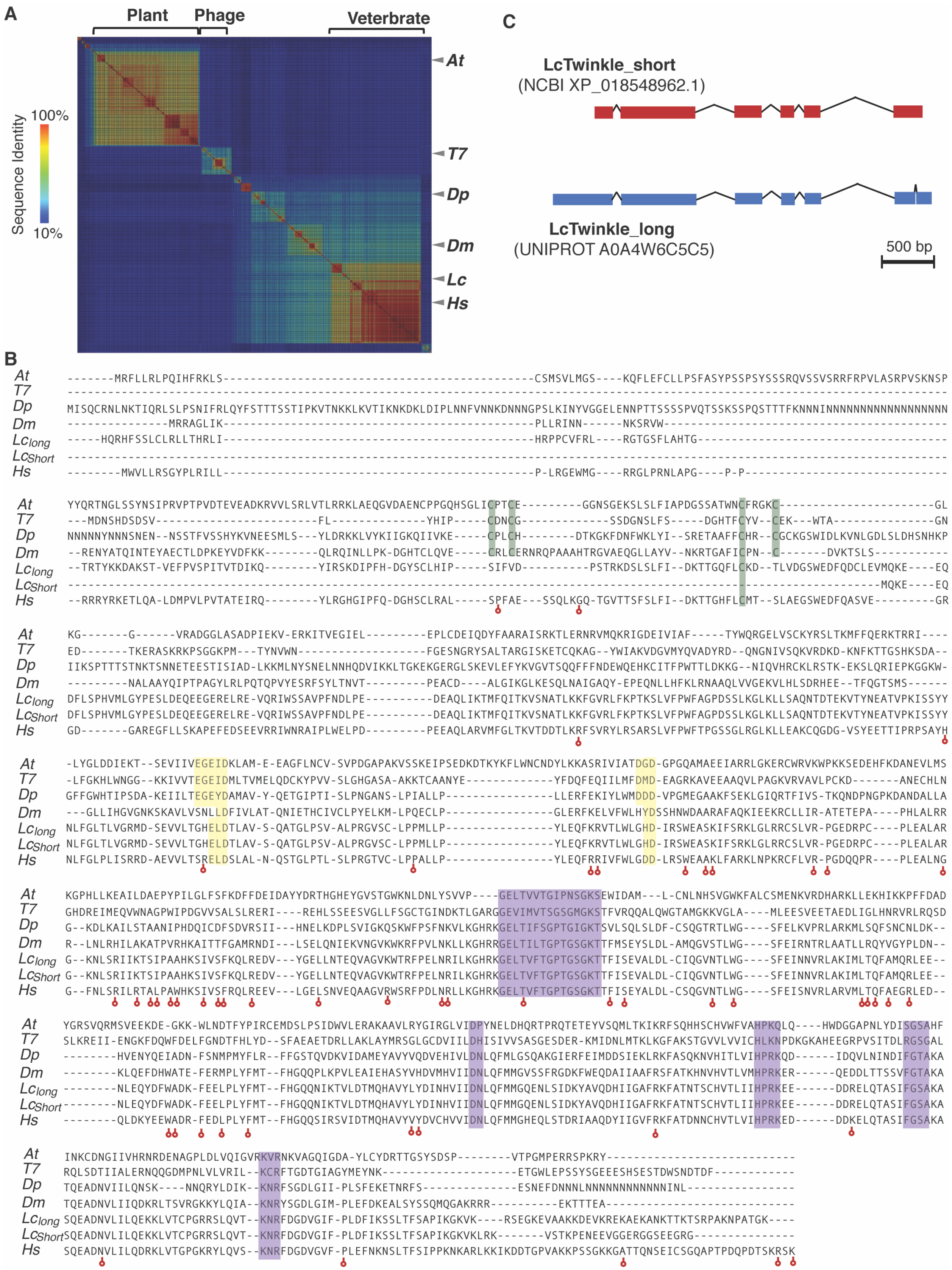
A
Plant
Phage
Veterbrate
Sequence Identity
100%
10%
At
T7
Dp
Dm
Lc
Hs
B
At
T7
Dp
Dm
Lclong
LcShort
Hs
C
LcTwinkle_short
(NCBI XP_018548962.1)
LcTwinkle_long
(UNIPROT A0A4W6C5C5)
500 bp

**Supplementary Figure 1. Sequence analysis of Twinkle helicase. A.** Sequence similarity matrix of Twinkle and its homologs. Each row or column in the matrix corresponds to a Twinkle homolog. The identity of the two sequences is color-coded according to the bar on the left. The positions of several representative Twinkle homologs are highlighted. *At, T7, Dp, Dm, Lc, Hs* are for *Arabidopsis thaliana, bacteriophage T7, Dictyostelium purpureum, Drosophila melanogaster, Lates calcarifer* and *homo sapiens,* respectively. *At*, *T7*, and *Dp* Twinkles have been reported to have primase activity. **B.** Sequence alignments of selected Twinkle homologs. The key residues for Zinc-binding domains, primase, and helicase active sites are colored in green, yellow and purple, respectively. The disease-related mutations are indicated by red arrows. **C.** Exons of the two variants of LcTwinkle. The short LcTwinkle is based on XP_018548962.1 from the NCBI protein database (https://www.ncbi.nlm.nih.gov/protein/), and the long LcTwinkle is from the uniport database (https://www.uniprot.org/) with access code A0A4W6C5C5.

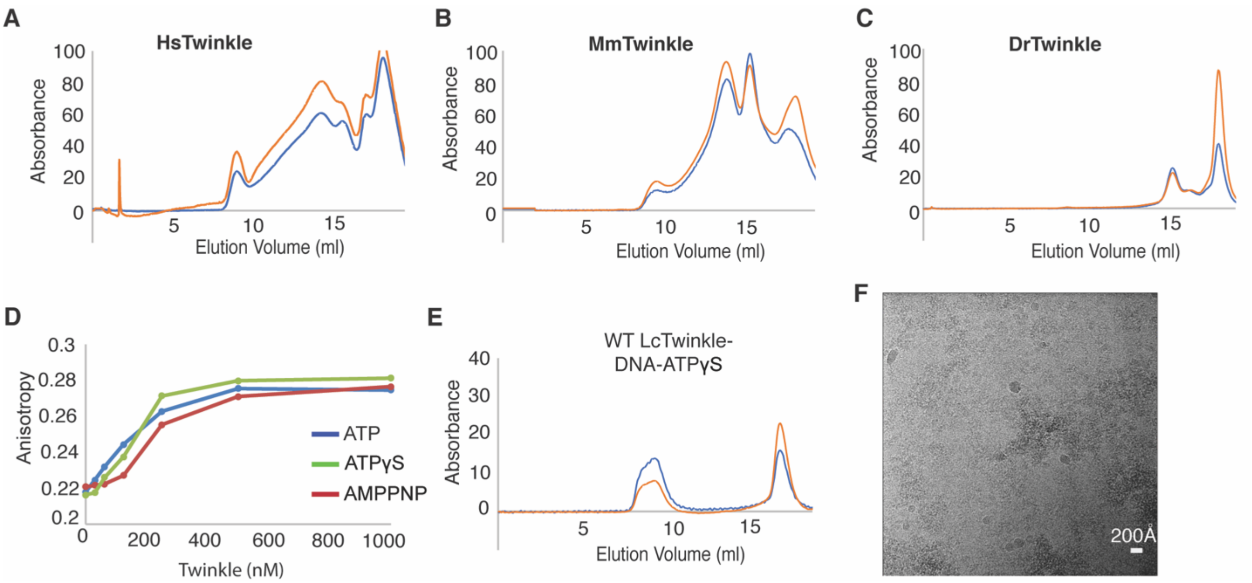

**Supplementary Figure 2. Characterization of Twinkle homologs and the WT LcTwinkle-DNA complex in the presence of ATPγS. A-C.** Gel filtration profile of Hs (**A**), Mm (**B**), and Dr (**C**)Twinkle with ATP and DNA. **D.** Fluorescence anisotropy DNA binding of the WT LcTwinkle in the presence of ATP and analogs. **E.** Gel filtration profile of LcTwinkle with ATP**γ**S and DNA. **F.** A representative cryo-EM micrograph of WT LcTwinkle complex with ATP**γ**S and DNA.

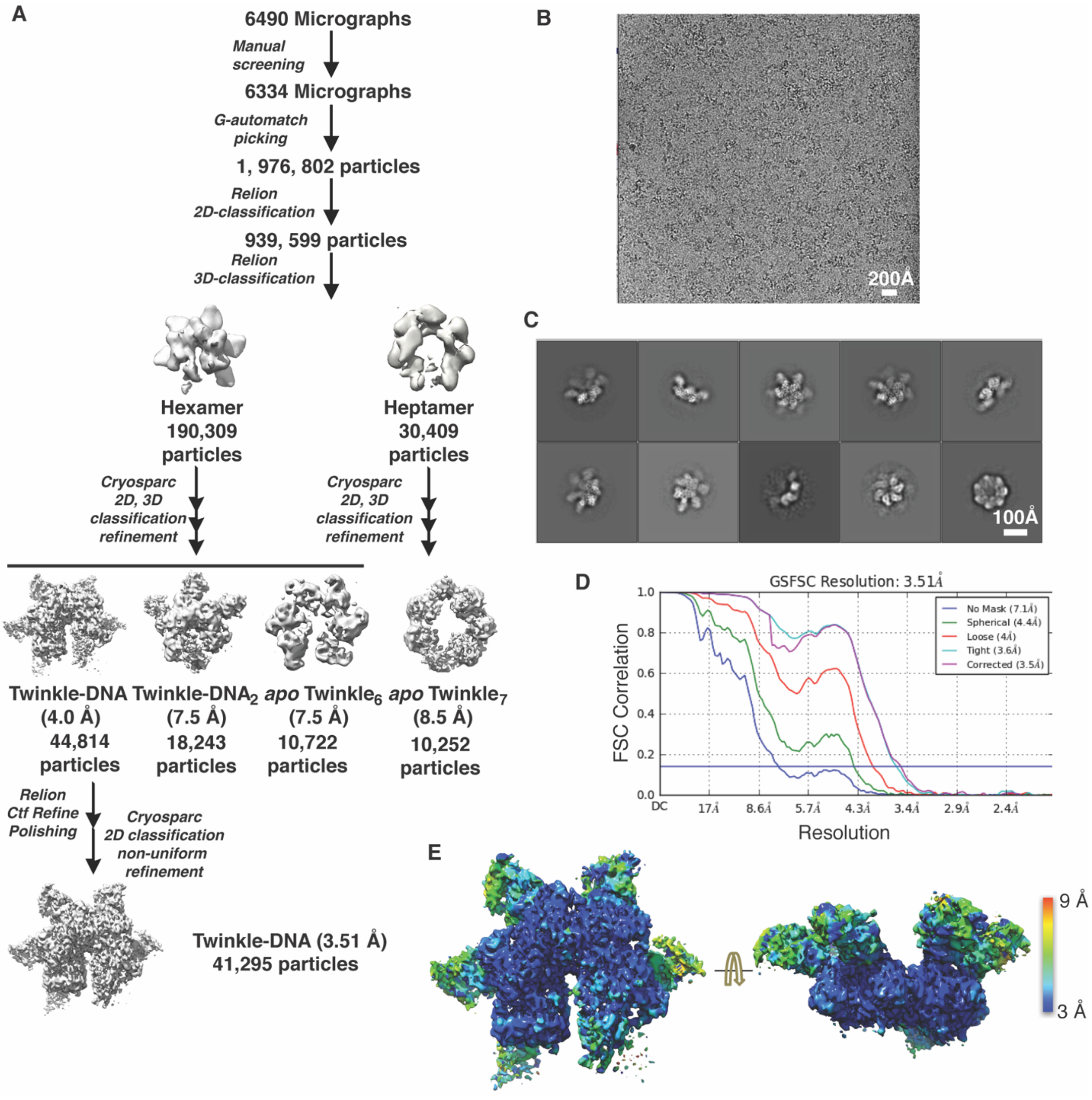

**Supplementary Figure 3. Cryo-EM data processing**. **A.** Overall data processing scheme. The software and number of particles at each step are indicated. **B.** A representative cryo-EM image of the LcTwinkle complex. **C.** 2D-classification averages. **D.** Fourier shell correlation (FSC) curve of the LcTwinkle-DNA complex. **E.** Local resolution estimation of the LcTwinkle-DNA complex. The local resolution is color-coded according to the color bar on the right.

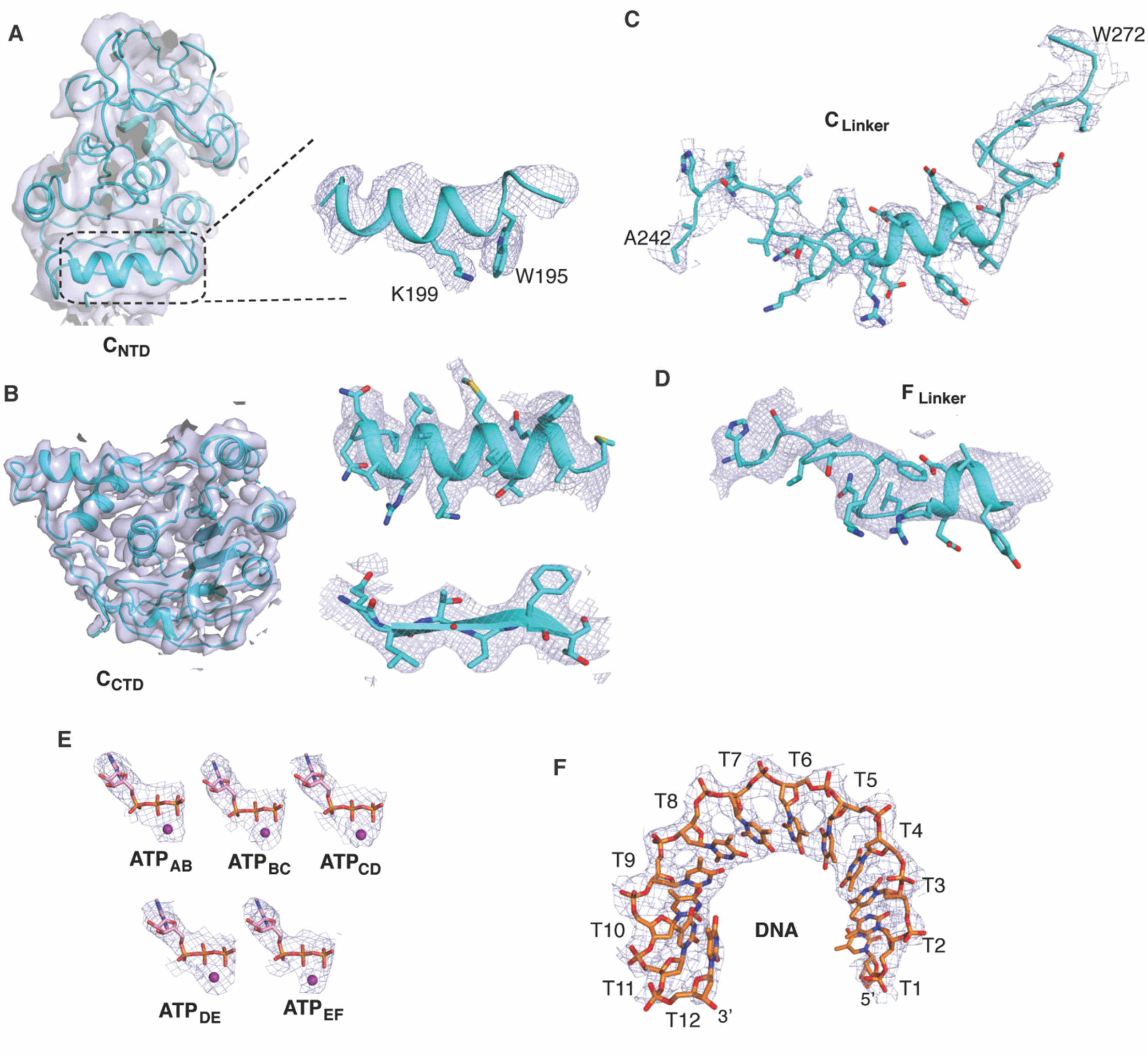

**Supplementary Figure 4. Local cryo-EM maps for the NTD (A), CTD (B), N-C linker of chain C (C), N-C linker of chain F (D), ATP (E), and DNA (F).**

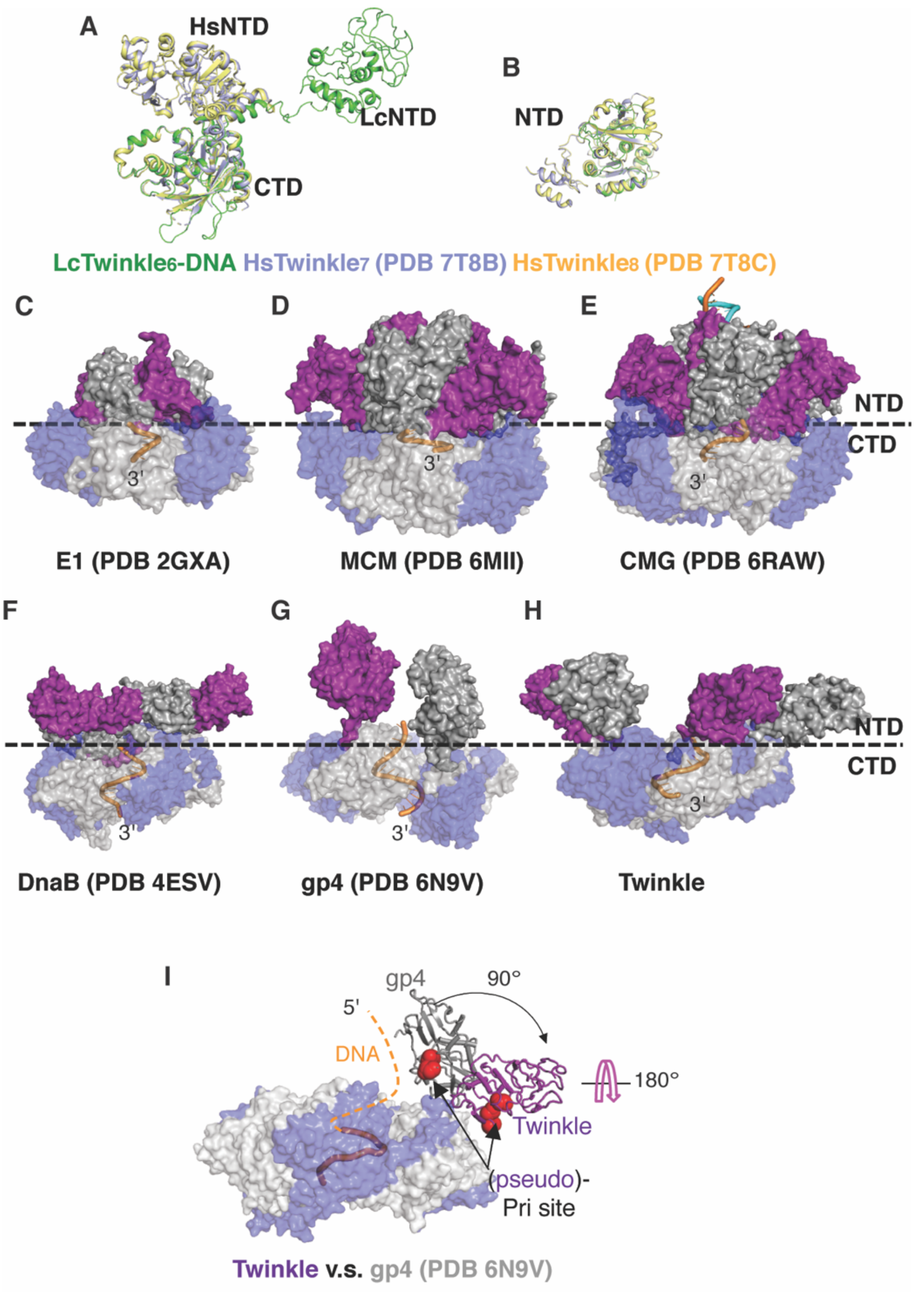

**Supplementary Figure 5. Unique conformation of LcTwinkle NTDs**. **A-B.** Structural comparison of the LcTwinkle with apo HsTwinkle with the CTD (**A**) or NTD (**B**) aligned. **C-H.** Structures of SF3 E1 (**C**), SF6 archaeal MCM (**D**), SF6 eukaryotic CMG (**E**), SF4 DnaB (**F**), SF4 gp4 (**G**), and SF4 Twinkle (**H**) helicases. In all panels, the NTDs are colored by alternative purple and dark grey. The CTDs are colored by alternative blue and light grey and set to be semi-transparent. The DNA is drawn as orange cartoon. **I.** Structural comparison of NTDs in gp4 and LcTwinkle. The primase active site (or pseudo active site) is highlighted by red spheres of the active site residues. The unwound DNA is indicated as dotted lines. The NTD rotation and flipping are marked by arrows.

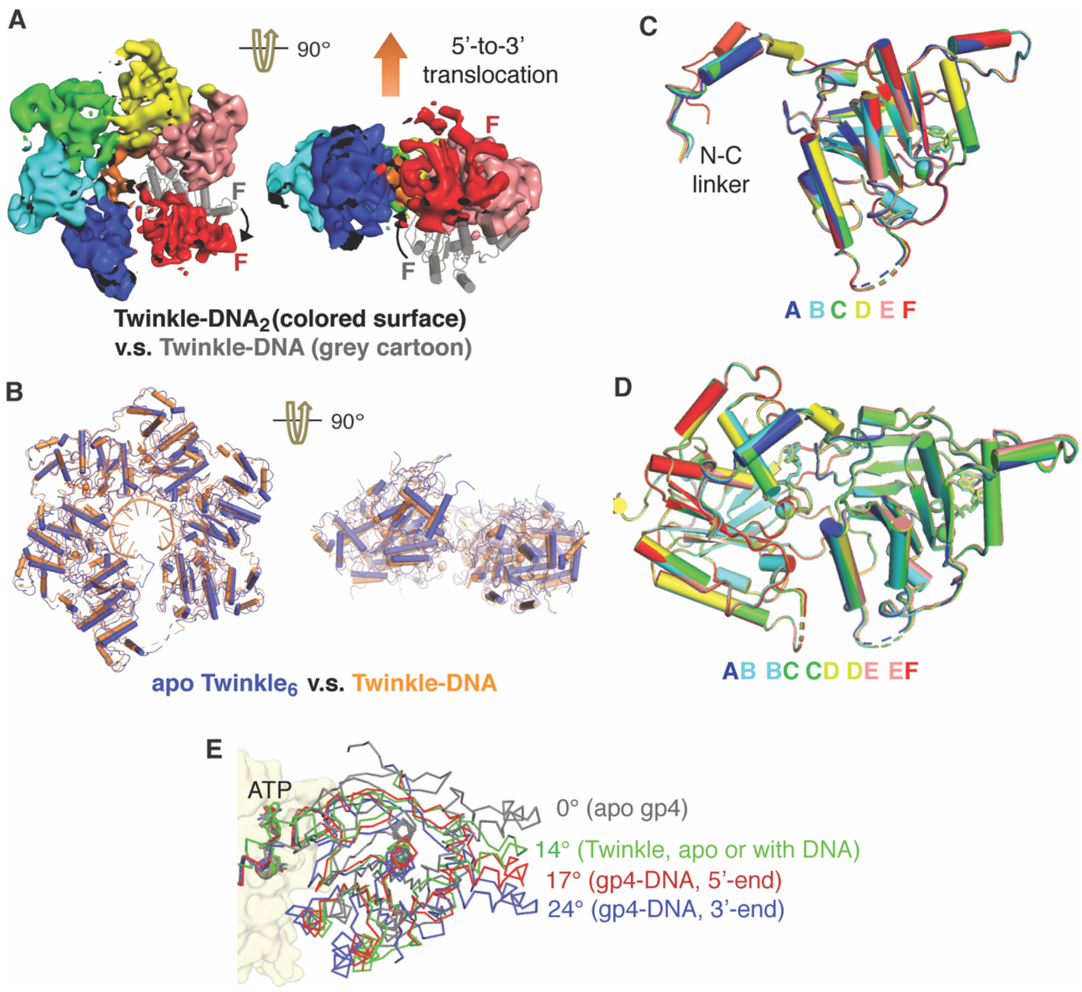

**Supplementary Figure 6. Helicase domain conformations in LcTwinkle structures. A.** Helicase domain movement in LcTwinkle-DNA$_2$. The movement of subunit F relative to that in the LcTwinkle-DNA complex is indicated by arrows. **B.** Structural comparison of LcTwinkle helicase domains in the presence and absence of DNA. **C.** Superposition of different subunits from LcTwinkle-DNA complexes suggests that they are superimposable. **D.** Superposition of different subunit-dimers from the LcTwinkle-DNA complex suggests that they are superimposable. **E.** Comparison of ATPase sites at the subunit-subunit interfaces in LcTwinkle-DNA and gp4-DNA complexes. The rotation angle relative to the planar apo gp4 dimer is indicated.

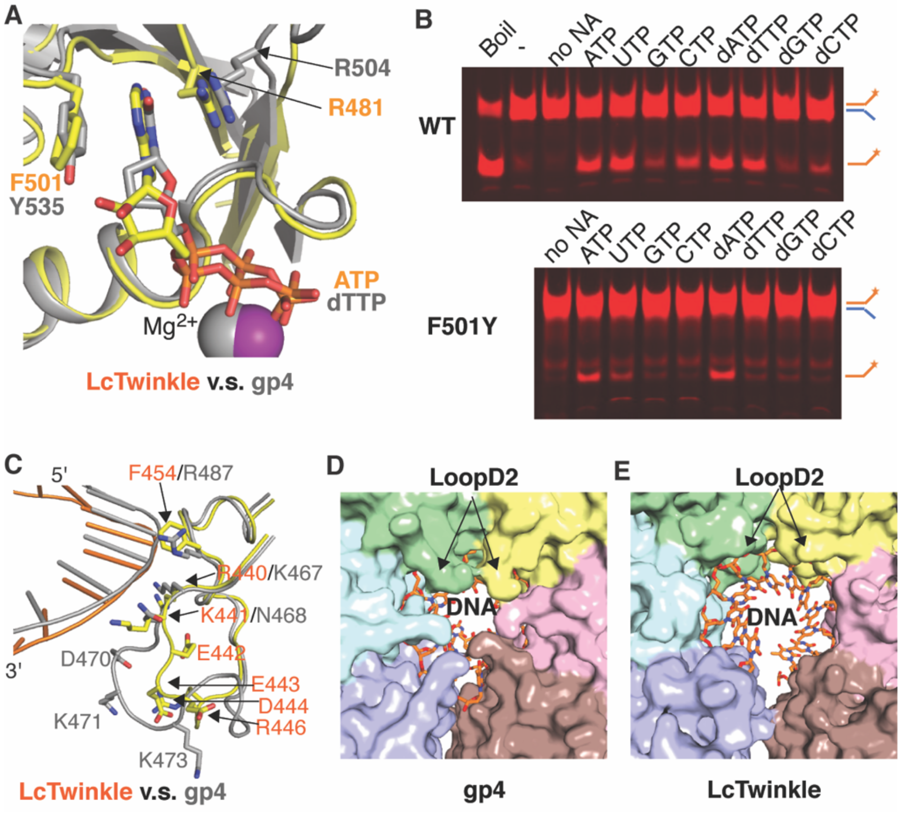

**Supplementary Figure 7. ATP and DNA binding in LcTwinkle. A.** Comparison of ATPase sites in LcTwinkle (colored) and gp4 (grey). The key residues for base recognition are labeled. **B.** Nucleotide triphosphate preference for WT and F501Y LcTwinkle. **C.** Structural alignment of DNA binding loops in LcTwinkle (colored) and gp4 (grey). **D-E.** View of DNA from the CTD side of the hexamer in gp4 (**D**) and LcTwinkle (**E**). The DNA is well-protected by LoopD2 in gp4 while the DNA in LcTwinkle is exposed.

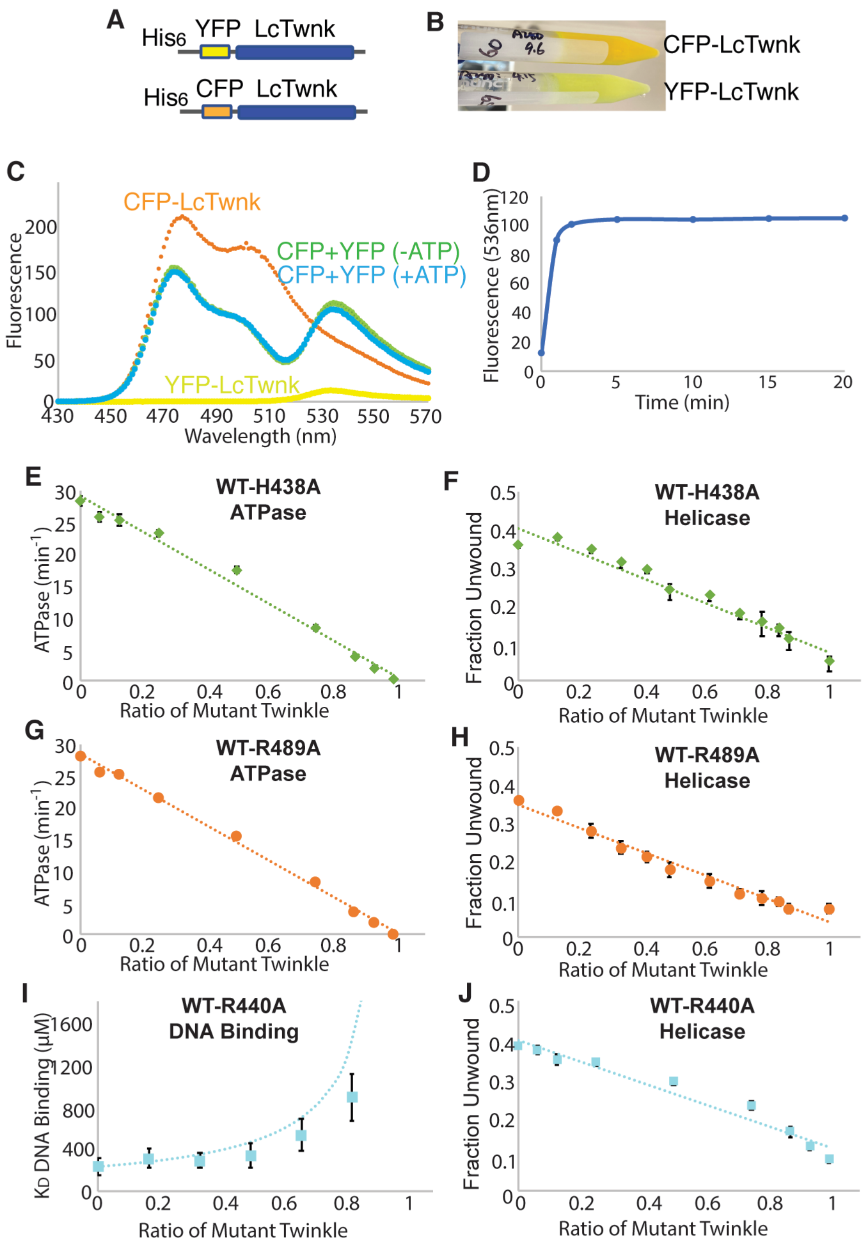

**Supplementary Figure 8. Additional subunit doping experiments. A.** contructs of CFP and YFP labeled Twinkle for confirming subunit exchange. **B.** The purified CFP and YFP labeled Twinkle. **C.** Fluorescence scans of CFP-Twinkle (orange), YFP-Twinkle (yellow), the mix of the two with (green) and without (blue) ATP. **D.** Time plot of the FRET peak at 536 nm suggest repaid subunit exchange of Twinkle. **E-J,** subunit doping experiments. Panels (**E**) and (**G**) are ATPase assays of WT with H438A (**E**) and R489A (**G**) LcTwinkle. Panels (**F**), (**H**), and (**J**) are helicase assays of WT with H438A (**F**), R489A (**H**), and R440A (**J**) LcTwinkle. Panel (**I**) is the

DNA binding assay of WT with R440A LcTwinkle. The error bars represent standard derivations from three independent experiments.

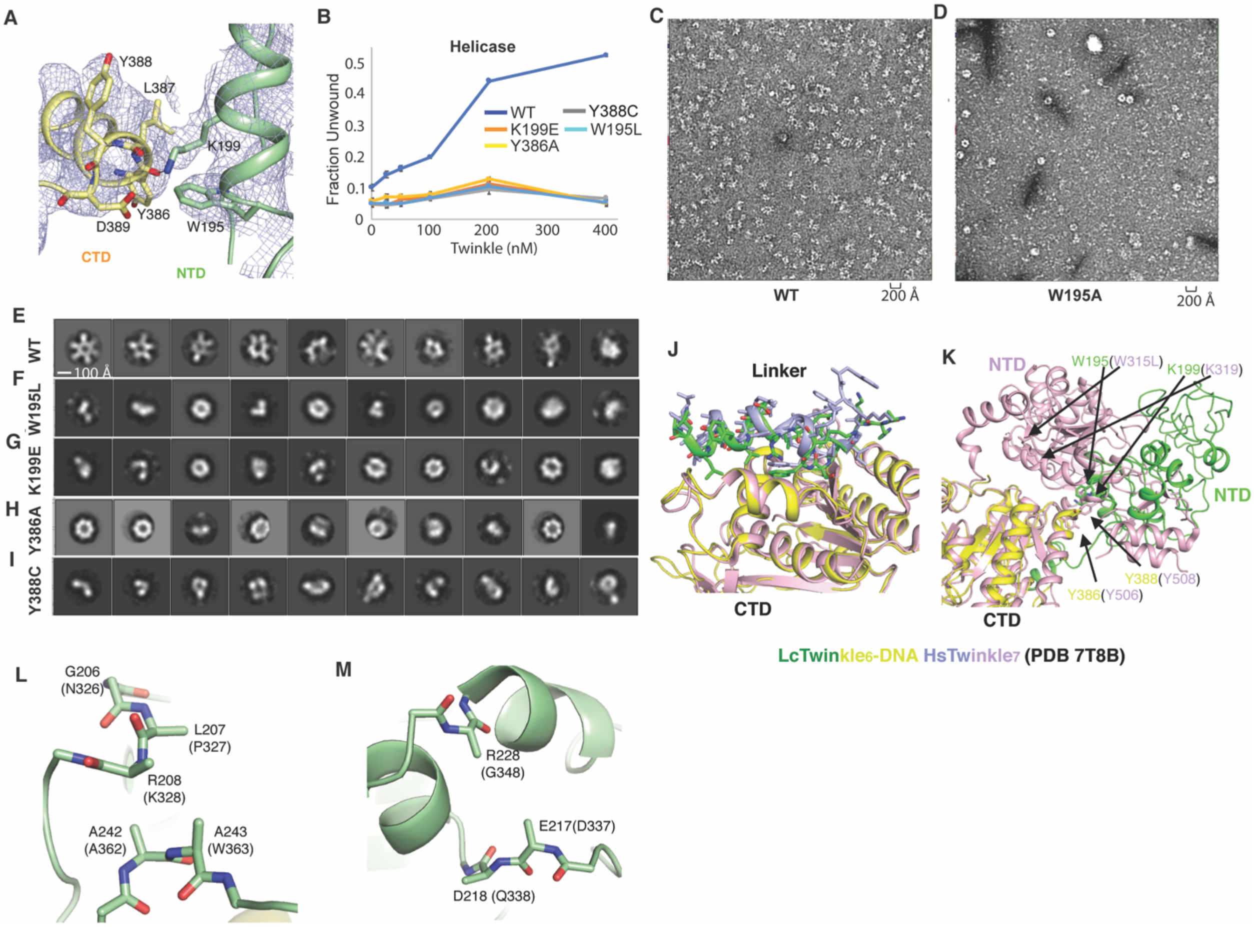

**Supplementary Figure 9. The NTD-CTD interface. A.** Structure of the NTD (green)-CTD (yellow) interface. The cryo-EM densities are shown as light blue mesh. **B.** Helicase activities of LcTwinkle with point mutations at the N-C interface. **C-D.** Representative negative staining images of WT (**C**) and W195A (**D**) LcTwinkle. **E-I.** 2D classes from negative staining EM of WT (**E**), W195L (**F**), K199E (**G**), Y386A (**H**) and Y388C (**I**) LcTwinkle. The class averages are ordered according to the number of particles in each class, with the first class with most particles. **J.** Structural comparison of the N-C linker interactions with the CTD in LcTwinkle-DNA complex and apo HsTwinkle. **K.** Structural comparison of the NTD-CTD interactions in LcTwinkle-DNA complex and apo HsTwinkle. **L.** The local environment of A243 (human Twinkle W363). **M.** The local environment of R228 (human Twinkle G348).

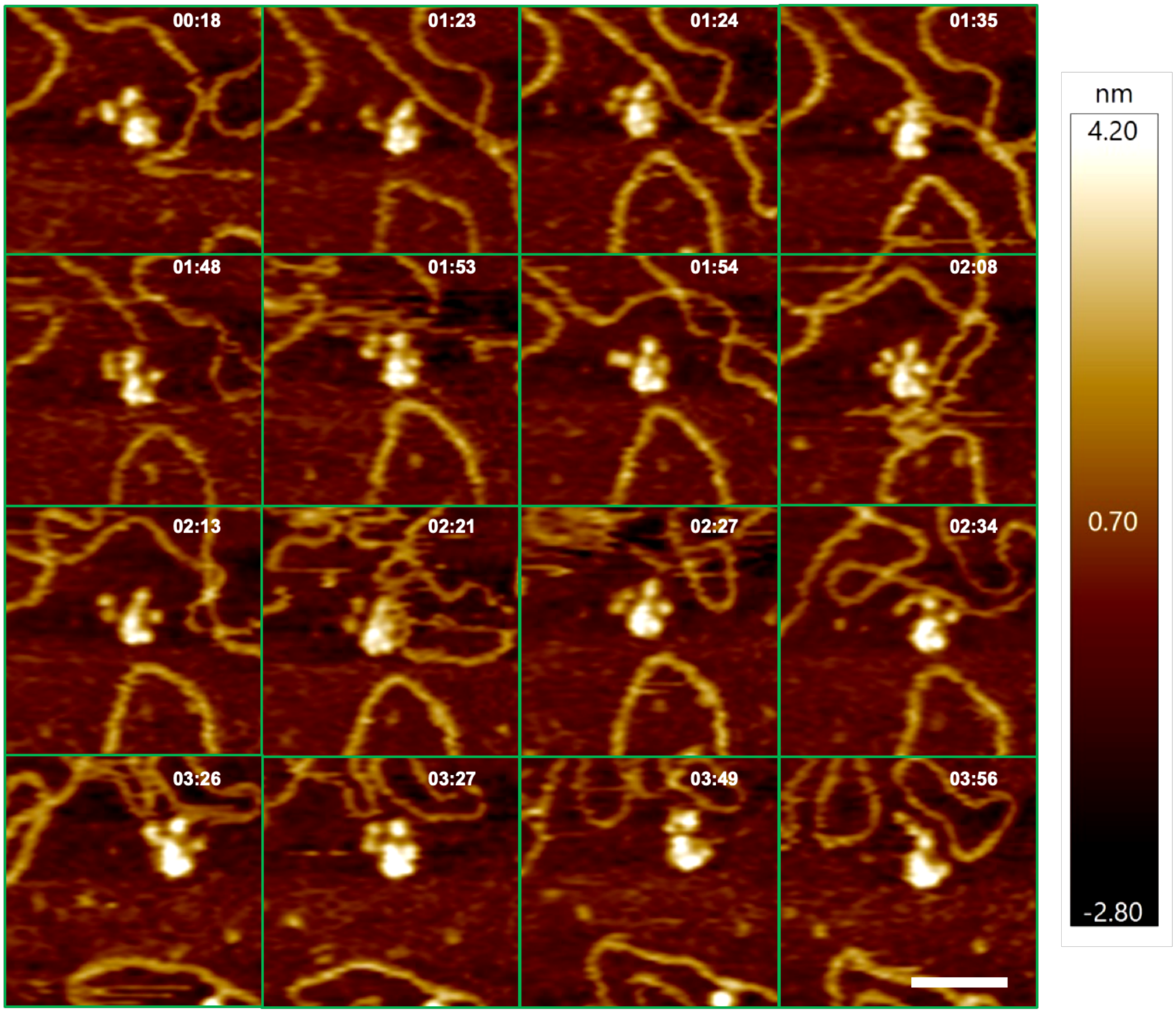

**Supplementary Figure 10. Snapshots from real-time HS-AFM imaging showing multiple N-protrusions on a single LcTwinkle molecule.** The LcTwinkle side with N-protrusions is highlighted by green arrows. Also see Video S4. XY Scale bar = 50 nm.

**Supplementary Table 1. Cryo-EM data collection, refinement and validation statistics**

| | LcTwinkle-DNA | LcTwinkle-$DNA_2$ | Apo $LcTwinkle_6$ | Apo LcTwinkle7 |
|---|---|---|---|---|
| PDB Code | | | | |
| EMDB Code | | | | |
| **Data collection and processing** | | | | |
| Magnification | 130,000 | 130,000 | 130,000 | 130,000 |
| Voltage (kV) | 300 | 300 | 300 | 300 |
| Electron exposure ($e^-/Å^2$) | 49 | 49 | 49 | 49 |
| Defocus range (μm) | 0.6-3 | 0.6-3 | 0.6-3 | 0.6-3 |
| Pixel size (Å) | 1.075 | 1.075 | 1.075 | 1.075 |
| Number of micrographs | | | | |
| Collected | 6490 | 6490 | 6490 | 6490 |
| Used | 6334 | 6334 | 6334 | 6334 |
| Symmetry imposed | C1 | C1 | C1 | C1 |
| No. particles used | 44814 | 18243 | 10722 | 10252 |
| Map resolution (Å) | | | | |
| Corrected, FSC=0.143 | 3.5 | 7.5 | 7.5 | 8.5 |
| **Refinement** | | | | |
| Map sharpening B ($Å^2$) | -110 | | | |
| Cross-correlation | | | | |
| CC_mask | 0.71 | | | |
| CC_volumes | 0.71 | | | |
| CC_peaks | 0.59 | | | |
| Model composition | | | | |
| Non-hydrogen atoms | 17092 | | | |
| Protein / DNA | 16932 | | | |
| Ligands / $Mg^{2+}$ | 160 | | | |

| | |
|---|---|
| B-factors | |
| Non-hydrogen atoms | 197.1 |
| Protein / DNA | 197.6 |
| Ligands / $Mg^{2+}$ | 145.0 |
| R.m.s deviations | |
| Bond lengths (Å) | 0.002 |
| Bond angles (º) | 0.558 |
| Ramachandran | |
| Favored (%) | 92.1 |
| Allowed (%) | 7.8 |
| Outlier (%) | 0.1 |
| Validation | |
| MolProbity score | 1.37 |
| Clashscore | 6.2 |
| Poor rotamers (%) | 0.4 |

**Supplementary Table 2. Steady-state kinetic analysis of Twinkle ATPase activity**

| | $K_M$ (µM) | $k_{cat}$ ($min^{-1}$) | $k_{cat}$ / $K_M$ ($min^{-1}.µM^{-1}$) |
|---|---|---|---|
| **WT** | 870±80 | 28±1 | 0.032 |
| **WT (no DNA)** | 930±90 | 18±1 | 0.019 |
| **E325Q** | 470±30 | 0.29±0.05 | 0.00055 |
| **R489A** | 580±50 | 1.7±0.4 | 0.0029 |
| **H438A** | 550±40 | 1.8±0.4 | 0.0033 |
| **S456A** | 760±70 | 15±1 | 0.020 |
| **R440A** | 670±80 | 15±1 | 0.022 |
| **W195L** | 7100±2000 | 55±15 | 0.0077 |
| **Y386A** | 8250±1800 | 59±13 | 0.0072 |
| **Y388C** | 5770±1000 | 47±10 | 0.0081 |
| **R481A** | 8340±2000 | 51±12 | 0.0061 |
| **F501A** | 8130±1500 | 42±10 | 0.0052 |
| **F501Y** | 7800±1200 | 55±10 | 0.0071 |

**Supplemental Video 1. A LcTwinkle molecule without DNA in proximity shows limited conformational changes**

**Supplemental Video 2. A LcTwinkle molecule with limited conformational changes away from DNA later displays N-protrusions towards nearby DNA (Related to Figure 4A)**

**Supplemental Video 3. A LcTwinkle molecule uses proximity DNA induced N-protrusion to capture nearby DNA (Related to Figure 4B)**

**Supplemental Video 4. A LcTwinkle molecule shows N-protrusion from multiple domains, aiming to capture nearby DNA (Related to Extended Data Figure 10)**

**Supplemental Video 5. A LcTwinkle molecule uses N-protrusion to capture nearby DNA that leads to a large conformational change of the whole protein (Related to Figure 4C)**

**Supplemental Video 6. Initial DNA binding by LcTwinkle through N-protrusion leads to subsequent DNA loading at the central channel (Related to Figure 4D).** Multiple DNA capture events at N-protrusion and the central channel and unloading from DNA were revealed in this video.

**Supplemental Video 7. Dynamics of DNA capture and release by a LcTwinkle molecule**
